# A parallel channel of state-dependent sensory signaling from the cholinergic basal forebrain to the auditory cortex

**DOI:** 10.1101/2022.05.05.490613

**Authors:** Fangchen Zhu, Sarah E. Elnozahy, Jennifer Lawlor, Kishore V. Kuchibhotla

## Abstract

Cholinergic basal forebrain (CBF) signaling exhibits multiple timescales of activity with classic, slow signals related to brain and behavioral states and faster, phasic signals reflecting behavioral events, including movement and reinforcement. Recent evidence suggests that the CBF may also exhibit fast, sensory-evoked responses. It remains unknown, however, whether such sensory signals target the sensory cortex and how they relate to local functional topography. Moreover, the extent to which fast and slow CBF activity interact has been largely unexplored. Here, we used simultaneous two-channel, two-photon imaging of CBF axons and auditory cortical (AC) neurons to reveal that CBF axons project a robust, non-habituating, and stimulus-specific sensory signal to the AC. Individual axon segments exhibited heterogeneous but stable tuning to auditory stimuli allowing stimulus identity to be decoded from the population. However, CBF axons displayed no tonotopy and their frequency tuning was uncoupled from that of nearby cortical neurons. Chemogenetic suppression revealed the auditory thalamus as a principal source of auditory information to the CBF. Finally, slow fluctuations in cholinergic activity modulated the fast, sensory-evoked signals in the same axons, suggesting that a multiplexed combination of fast and slow signals is projected from the CBF to the AC. Taken together, our work demonstrates a novel, non-canonical function of the CBF as a parallel channel of state-dependent sensory signaling to the sensory cortex that provides repeated representations of a broad range of sound stimuli at all points on the tonotopic map.

## Introduction

The cholinergic basal forebrain (CBF) is the primary source of acetylcholine to the neocortex, hippocampus, and amygdala^1–5^. CBF signals are implicated in modulating attention^6–10^, supporting memory encoding^11–15^, and shaping cortical plasticity^16–20^. However, the classic view of cholinergic neuromodulation as slow, spatially diffuse, and regionally non-specific is rapidly evolving^21–23^. Anatomical studies have revealed a more structured organization of projections from the CBF^4,5,24–27^ and behavioral studies indicate that cholinergic neuromodulation operates at multiple timescales to convey different facets of information – slower tonic signals reflect modulations in internal state and behavioral contexts^28–33^ while faster phasic signals are associated with reinforcement^34–37^, movement^35,38–40^, and even sensory cues^41,42^. Fast CBF transients that are regionally-specific and tied to environmental features may provide a complement to slower, diffuse signaling of brain state in influencing downstream cortical networks. In particular, native cholinergic activity in response to neutral sensory cues has previously been observed using bulk calcium photometry in the basal forebrain^41,42^, suggesting that CBF may relay sensory information to downstream regions. However, it remains unknown whether such rapid sensory signaling target sensory cortices, and how it relates to the local functional topography. Moreover, little is known about the interactions between signaling at different timescales by the cholinergic system. Here, we used two-color, two-photon microscopy to record the activity of CBF axons and cortical neurons in the auditory cortex to investigate the spatiotemporal characteristics of sensory-evoked cholinergic activity.

## Results

### Cholinergic neuromodulation relays sensory information about neutral auditory stimuli to auditory cortex

CBF neurons in the basal forebrain have previously been observed to respond to auditory stimuli^41,42^. We investigated the extent to which cholinergic signals relay auditory information to the auditory cortex – a downstream cortical target, using two-photon microscopy to record the activity of CBF axonal projections to the auditory cortex. We expressed an axon-targeted variant of the genetically encoded calcium indicator GCaMP6s (axon-GCaMP6s), specifically in cholinergic neurons using a cre-dependent viral injection in the basal forebrain of ChAT-cre mice and recorded the calcium activity of CBF axonal projections to the auditory cortex (n = 8; **Fig. 1a-b**, **Supplementary Fig. 1**). Our optical approach allowed us to investigate both the spatial and temporal dynamics of cholinergic signals in subcellular axonal processes (**Fig. 1c**, example animal). In total, we identified 15,777 CBF axonal segments in 73 sites across the auditory cortex of 8 animals (n = 9±7 sites per animal). We presented passively-listening head-fixed animals with 20 repetitions of a white noise stimulus (100ms, 70-80 dB SPL) and observed multiple axonal segments that were significantly responsive to the neutral stimulus (**Fig. 1d-f**). Across 8 animals, 24.8±21.9% of identified axon segments responded to white noise and were distributed across the auditory cortex (**Fig. 1g**, example animal, **Supplementary Fig. 2**). We observed that a similar percentage of axon segments responded to frequency up-sweeps (24.6±18.8%) and down-sweeps (22.3±11.8%) across the broad extent of the auditory cortex (**Supplementary Fig. 2**).

**Figure 1.**
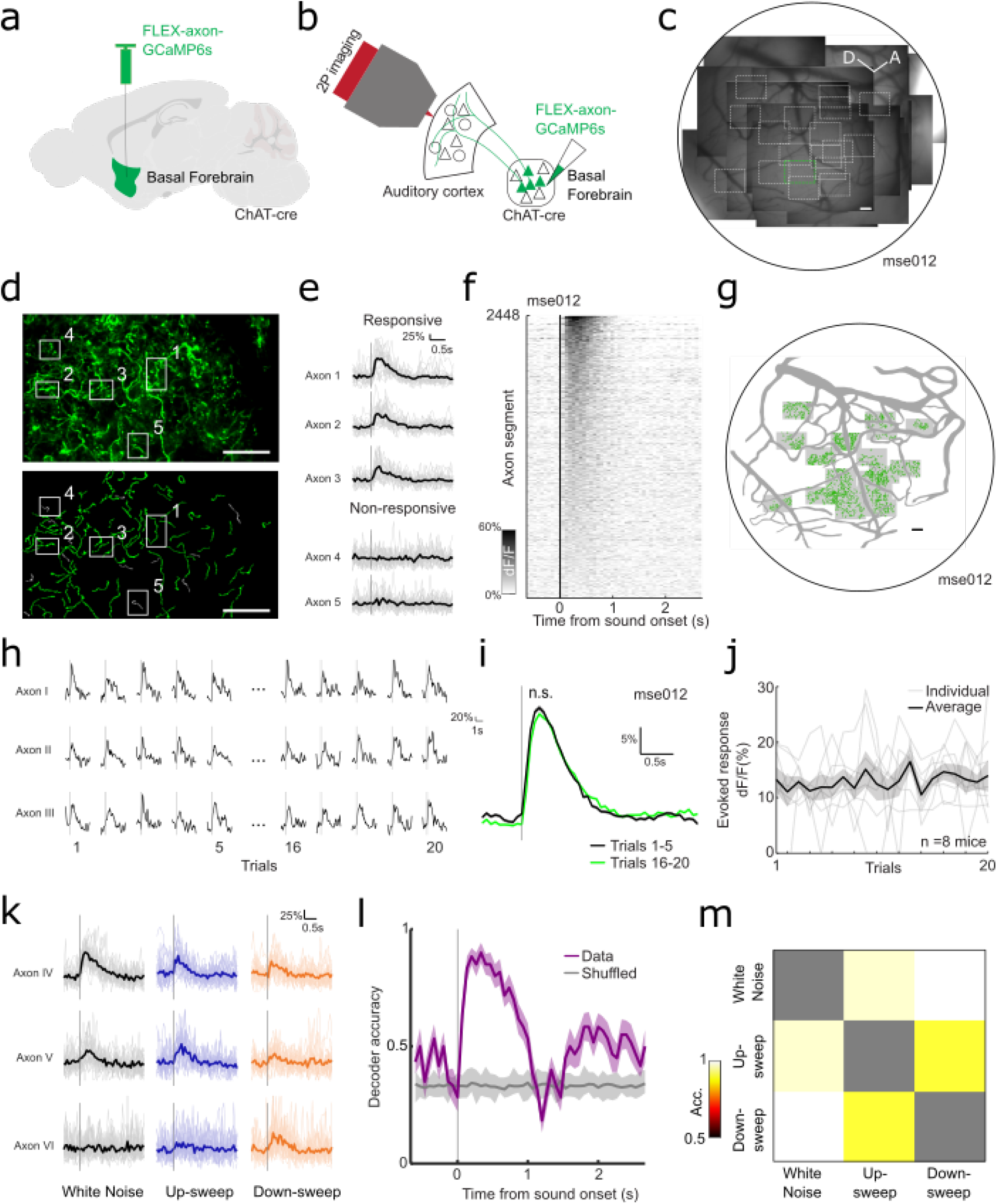
Robust, non-habituating, and stimulus-specific auditory response of cholinergic axons. (**a**) Schematic of basal forebrain viral injection. (**b**) Schematic of CBF projection to auditory cortex and imaging above auditory cortex. (**c**) Composite widefield image of all recording sites in one example animal. Black border demarcates approximate location of cranial window and white boxes indicate two-photon imaging sites at 4x magnification. Green box indicates location of example site in (**d**). Scalebar = 100μm (**d**) Top: Mean fluorescence image of cholinergic axons (green, axon-GCaMP6s) in example recording site. Bottom: manually identified axon ROIs of example site. Responsivity of example axon ROIs in boxes 1-5 are shown in (**e**). Scalebar = 50μm (**e**) Example traces of axon ROIs that are responsive and non-responsive to white noise presentation. Bold line indicates mean response across 20 presentations, faded traces indicate individual presentations of white noise. Gray lines indicate presentation of white noise. (**f**) Heatmap of average evoked response (ΔF/F) to white noise for all identified axon segments in one animal (n = 2448 axon segments). (**g**) Spatial distribution of axon segments responsive to white noise (green) in one animal. Shaded boxes indicate recording sites. Scalebar = 100μm (**h**) Fluorescence trace of example axon ROIs for 1-5 and 16-20 presentation of white-noise stimulus. Gray lines indicate presentation of white noise. (**i**) Mean fluorescence trace of all axon ROIs in one example animal for 1-5 (black) and 16-20 (green) presentation of white noise stimulus, p = 0.412. Gray line indicates presentation of white noise and shaded region indicates SEM. (**j**) Amplitude of evoked response for white noise across 20 presentations for all animals (n = 8 animals). (**k**) Example traces of axon ROIs that are responsive to white noise, up-sweeps and down-sweeps. Bold line indicates mean response across 20 presentations, faded traces indicate individual presentations of white noise. Gray lines indicate presentation of auditory stimulus. (**l**) Decoding accuracy of multi-class decoder predicting the identity of auditory stimuli from population axonal activity (white noise, up- and down-sweeps). (**m**) Pairwise population decoding of white noise, up-sweep and down-sweep.

To determine whether the cholinergic transients are sensory responses, we investigated a few alterative explanations. It is possible that these robust transients indicate the detection of novel, unexpected stimuli^42,43^. If so, we would expect substantial habituation after repeated presentations of the same stimulus. We compared the mean response amplitude of the first five presentations of white noise to that of the last five presentation and found no significant difference (p = 0.412; **Fig. 1h-i**). Across the 20 presentations of the stimulus, the mean amplitude of the evoked response remained relatively constant, indicating a non-habituating response that is not only driven by novelty (**Fig. 1j**). Another possibility is that the phasic transients arise due to micro-movements of the animal when the auditory stimuli are detected^35^,^38–40^. We extracted the precise timing of movements during the recording sessions and found that 81.6% of the evoked signals were not associated with micro-movements (**Supplementary Fig. 3**). Cholinergic axons thus exhibit non-habituating phasic transient that is time-locked to stimulus-presentation, all of which are hallmarks of sensory responses.

We further observed that CBF axons displayed different degrees of responsivity to the complex sounds presented (**Fig. 1k**). Hence, we asked if the cholinergic signals can do more than just convey the detection of an auditory stimulus and instead play a direct sensory role relaying information about stimulus identity to the auditory cortex. To test this, we trained a linear decoder to predict the identity of the complex sound stimuli (white noise, up-sweep, or down-sweep) from the population activity of all axons. We observed high accuracy of sound-identity decoding well above 80% (chance level = 33.3%) after sound presentation suggesting that the cholinergic signal is stimulus-specific (**Fig. 1l**). To further investigate if the decoding is driven by specific stimuli, we tested each pair of complex sounds and observed robust pairwise decoding suggesting that phasic, cholinergic neuromodulation carries identifying information about individual auditory stimulus (**Fig. 1m**). Robust stimulus-identity decoding was also evident within individual animals (**Supplementary Fig. 4**). Taken together, our data argue that the CBF provides a parallel pathway for sensory signals of neutral auditory stimuli to the auditory cortex.

### Cholinergic axons display heterogeneous frequency-specific response to pure tones

The central auditory system exhibits a precise topography of frequency coding (tonotopy) that begins in the cochlea and propagates through the feedforward hierarchy to the auditory cortex. Having demonstrated that cholinergic signals also relay auditory information to the auditory cortex, we asked whether CBF axons exhibit frequency tuning. We presented half-octave spaced pure tone stimuli in a pseudorandom order to passively listening animals and recorded sound-evoked phasic responses from individual cholinergic axon segments (n = 15,777). We observed that CBF axons displayed frequency tuning - axon segments responded robustly and reliably to particular frequencies and the response amplitude decreased for frequencies further away from their best frequency (**Fig. 2a-b**). Furthermore, CBF axons exhibited a broad range of frequency responsivity: 82.7% of all identified axon segments responded to 1-2 of the presented pure tones, while 0.6% responded to 5-6 tones (**Fig. 2c**). Notably, more axon segments responded to the frequencies between 4.8kHz to 19kHz compared to frequencies above 19kHz (**Fig. 2d**).

**Figure 2.**
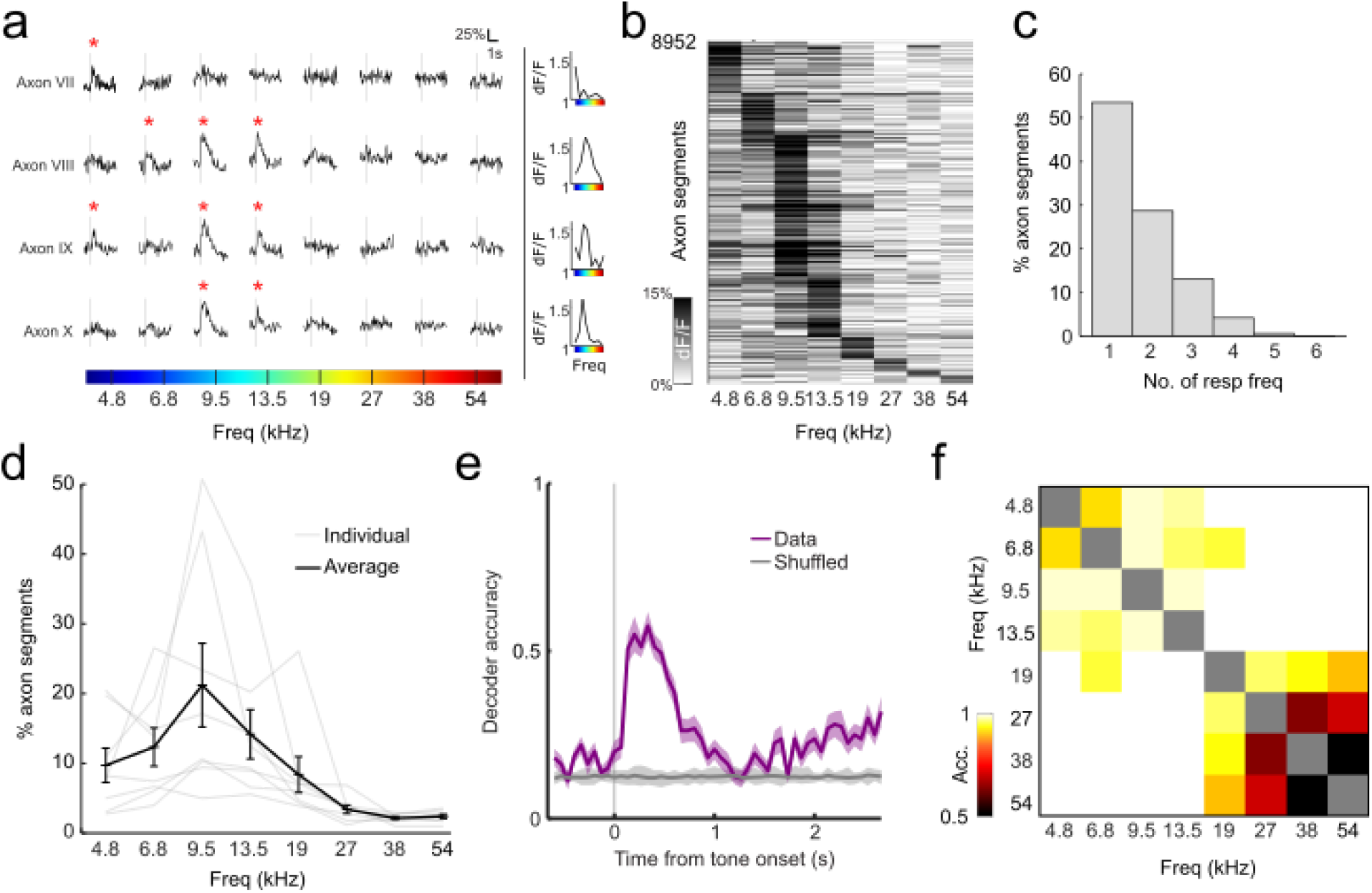
Frequency-specific tuning of cholinergic axons. (**a**) Selective evoked responses to pure tones in 4 example axon segments. Gray lines indicate presentation of auditory stimulus and red asterisks indicate significant responses. Tuning curve for each axon is plotted on the right. (**b**) Heatmap of amplitude of evoked response to pure tones in responsive axons. (**c**) Proportion of responsive axon segments that respond to various numbers of pure tones (**d**) Proportion of sound-responsive axon segments that responded to each pure tone for all animals (n = 8 animals). (**e**) Decoding accuracy of multi-class decoder predicting the identity of pure tone presented from population axonal activity. (**f**) Pairwise population decoding of 8 pure tones presented.

Given the observed heterogeneity in CBF axonal responses to pure tones, we asked whether cholinergic signals carried information about the frequency of auditory stimuli. Using the similar approach described above, we trained a multi-class decoder on the eight pure tones and found that tone identity could be decoded well above 50% accuracy (chance level = 12.5%) from population activity after tone presentation (**Fig. 2e**). Pairwise decoding of all stimuli pairs revealed that there is robust pairwise decoding for tones in the low-mid frequency of the mice hearing range suggesting that cholinergic transients carry information about those frequencies (**Fig. 2f**). Robust stimulus-identity decoding was also evident in individual animals (**Supplementary Fig. 5**). Taken together, our results argue that cholinergic axons display tuning properties that allow it to project a frequency-specific representation of auditory stimuli to the auditory cortex.

### CBF axons provides repeated representations of a broad range of frequencies at all points on the tonotopic map

Frequency-specific responses of CBF axons give rise to the possibility of a finer topography of functional cholinergic activity in the tonotopically-organized auditory cortex. Auditory cortical neurons display a tonotopy along the rostro-caudal axis^44,45^ which presents a powerful basis to compare the organizational specificity of functional cholinergic tuning. We used two-color, two-photon microscopy of CBF axons and cortical neurons to investigate whether the frequency tuning of cholinergic projections to the auditory cortex displayed any spatial organization and the relation between cholinergic tuning and the underlying cortical tonotopy. First, we expressed axon-GCaMP6s in CBF neurons of ChAT-cre mice that also expressed the red fluorescent calcium indicator, jRGECO1a, in auditory cortical neurons (see Methods). Using two-photon microscopy, we identified cholinergic axon segments (green, axon-GCaMP6s) innervating the primary auditory cortex (red, jRGECO1a) (**Fig. 3a-b**, example animal). We quantified the change in best frequency of these axon segments and observed no significant changes along the rostro-caudal axis (**Fig. 3c-d**, example site). This is in stark contrast with the striking tonotopic gradient found in cortical neurons in the primary auditory cortex recorded in animals expressing a similar calcium indicator (GCaMP6f) in auditory cortical neurons (**Fig. 3e**, **Supplementary Fig. 6**). These data suggest that cholinergic axons display minimal tonotopy compared to cortical neurons in the primary auditory cortex.

**Figure 3.**
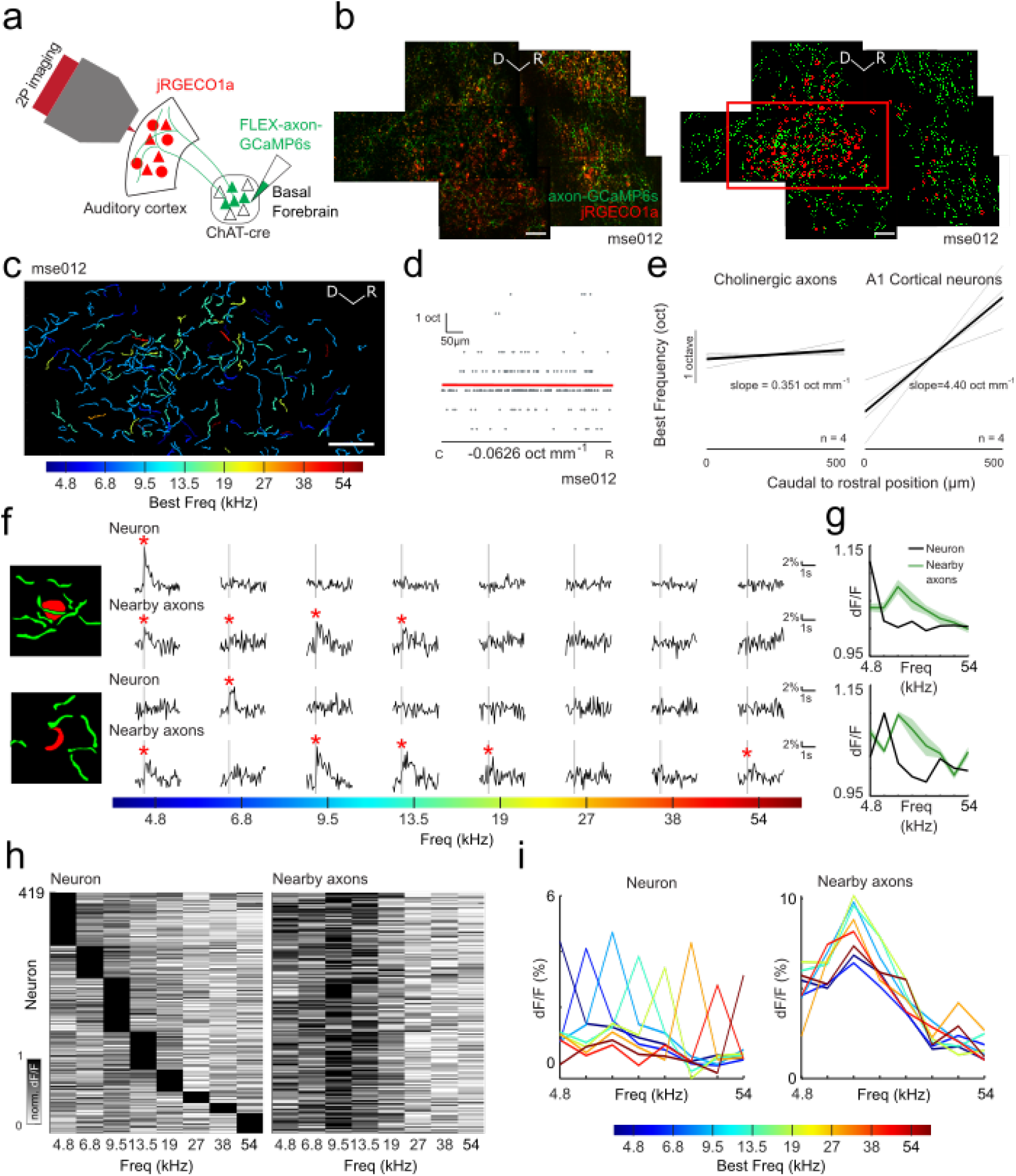
Frequency tuning of cholinergic axons uncoupled from tuning of cortical neurons. (**a**) Schematic of CBF projection to auditory cortex showing imaging strategy. (**b**) Left: mean composite fluorescence image of cholinergic axons (green) and cortical neurons (red) in example animal. Right: manually identified axon (green) and neuron (red) ROIs. Only responsive ROIs are shown. Red box indicates location of field of view in (**c**). Scalebar = 50μm (**c**) Axon ROIs colored by their best frequencies. Scalebar = 50μm (**d**) Change in best frequency of axon ROIs in (**c**) along the caudal-rostral axis. Scalebar = 50μm (**e**) Comparison of average change in best frequency for axon ROIs (n = 4 sites) and neuron ROIs in primary auditory cortex (n = 4 sites). (**f**) Left: schematic of example neurons and nearby axon segments (within 20μm). Right: mean evoked response of neuron and nearby axon segments to pure tone stimuli. Gray lines indicate presentation of auditory stimulus (**g**) Frequency tuning curve of example neurons (black) and nearby axon segments (green) in (**f**). (**h**) Left: normalized evoked response to pure tones of cortical neurons (n = 419 neurons). Right: normalized mean evoked response to pure tones of the nearby axon segments of the neuron in the corresponding row of the left heatmap. (**i**) Left: mean tuning curve of cortical neurons grouped by their best frequency. Right: mean tuning curve of the nearby axon segments of cortical neurons grouped by best frequency of cortical neurons.

However, it is possible that the responses of local axonal segments may overlap with the preferred frequencies of adjacent auditory cortical neurons. Hence, we compared the tuning of auditory cortical neurons and their nearby cholinergic axons directly. We identified 419 tone-responsive cortical neurons and their respective nearby axon segments in 6 animals (**Fig. 3b**, example animal). We found many single-peak neurons that were tuned to particular frequencies as expected (**Fig. 3f-g**). Interestingly, local axon segments were not co-tuned with the cortical neuron (**Fig. 3f-g**), but were instead responsive to a wider range of frequencies (**Fig. 3h**). When we compared the tuning profile of all the auditory cortical neurons with their nearby axons, we observed that, regardless of the tuning of the cortical neuron, the local cholinergic axon segments responded most to frequencies between 4.8kHz to 19kHz (**Fig. 3i**), whereas the local cortical neurons tuning was more similar (**Supplementary Fig. 7**). These data reveal that the sensory information relayed by CBF axons are largely uncoupled from cortical neuronal tuning, thereby providing a scaffold for interaction between parallel streams of sensory information to the auditory cortex.

### The medial geniculate body sends auditory information to the cholinergic basal forebrain

Our findings that cholinergic axons relay auditory information to the cortex raise the question of where along the ascending auditory pathway is the source of auditory information to the CBF. Previous anatomical studies have revealed that the CBF receives dense innervations from the medial geniculate body in the thalamus (‘auditory thalamus’)^3,46^. We investigated whether the auditory thalamus relays auditory information to the CBF. We performed chemogenetic suppression of the auditory thalamus using inhibitory designer receptors exclusively active by designer drugs (DREADDs) hM4Di and examined its effect on the tuning response of cholinergic axons in the auditory cortex (**Fig. 4a**, **Supplementary Fig. 8**). Consistent with the findings above, cholinergic projections to the auditory cortex in these animals displayed robust evoked responses to pure-tones (**Fig. 4b-c**). Intraperitoneal injection of clozapine N-oxide (CNO) suppressed activity in the medial geniculate body, which we confirmed by observing attenuated sound-evoked responses in cortical neurons (**Supplementary Fig. 9**). MGB suppression resulted in marked reduction of percentage of responsive CBF axons (after saline injection: 59.9±11.2%, after CNO injection: 37.3±18.0%, p<0.05) and a significant attenuation of sound-evoked CBF axonal responses (F(1,48) = 27.67, p<0.001); **Fig. 4b-d**).

**Figure 4.**
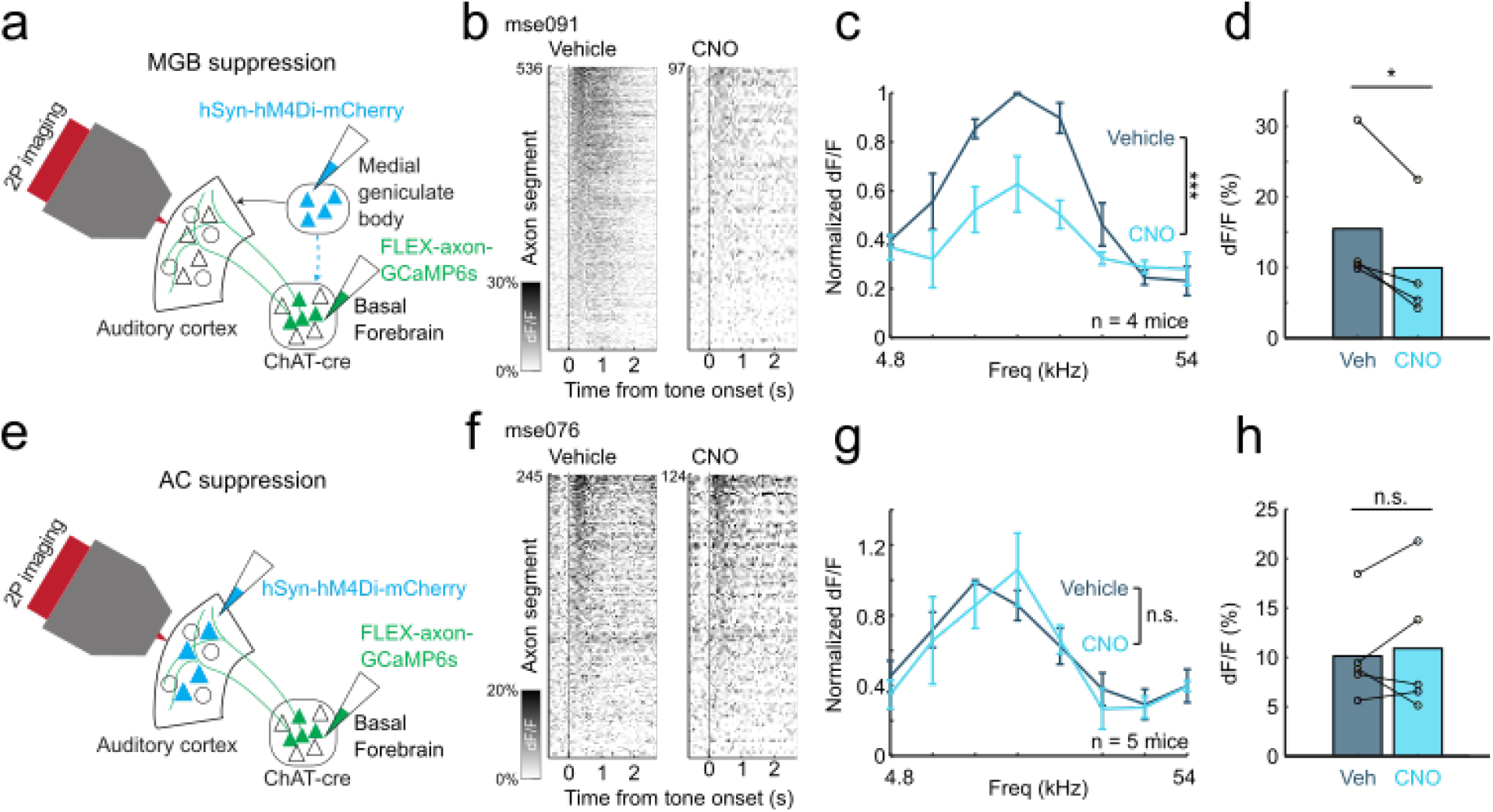
Suppression of auditory thalamus but not auditory cortex attenuates sound-evoked cholinergic responses. (**a**) Schematic of injection strategy for suppression of the medial geniculate body. (**b**) Evoked response in cholinergic axon segments to most responsive frequencies (9.5-19kHz) after intraperitoneal saline (left) and CNO injection (right) for an example animal. (**c**) Normalized evoked response to pure tones after intraperitoneal saline and CNO injection (n = 4 animals). Evoked response is significantly attenuated after CNO injection F(1,48) = 27.67, p<0.001. (**d**) Mean evoked response to most responsive frequencies (p<0.05; n = 4 animals). (**e**) Schematic of injection strategy for suppression of the auditory cortex. (**f**) Evoked response in cholinergic axon segments to most responsive frequencies (9.5-19kHz) after intraperitoneal saline (left) and CNO injection (right) for an example animal. (**g**) Normalized evoked response to pure tones after intraperitoneal saline and CNO injection (n = 5 animals). Evoked response is not attenuated after CNO injection, F(1,64) = 0.01, p = 0.908. (**h**) Mean evoked response to most responsive frequencies (p = 0.76; n = 5 animals).

It is also possible that the auditory thalamus relays information to the basal forebrain through the auditory cortex. To test that possibility, we chemogenetically suppressed the auditory cortex while recording cholinergic axonal response to pure tones (**Fig. 4e, Supplementary Fig. 8**). Intraperitoneal injection of CNO attenuated sound-evoked responses in auditory cortical neurons (**Supplementary Fig. 9**) but did not affect percentage of responsive CBF axons (after saline injection: 50.5±16.8%, after CNO injection: 50.0±35.6%, p = 0.958) or sound-evoked responses of CBF axons (F(1,64) = 0.01, p = 0.908) suggesting that the auditory cortex plays a minimal role in auditory information relay to the basal forebrain (**Fig. 4f-h**). These data together point to the auditory thalamus as a primary source of auditory input to the CBF.

### Tonic state-dependent cholinergic activity modulates phasic responses

The classic view of cholinergic neuromodulation proposes that the slow, diffuse signals from the CBF is a reflection of brain and behavioral states^28–33^. However, it is unknown how these tonic signals affect phasic transients from the same cholinergic neurons. We investigated the relation between phasic sensory-evoked responses and tonic state-dependent activity from the CBF using our optical approach which allowed us to detect changes in cholinergic activity at multiple timescales. During our recordings, we observed large endogenous fluctuations of baseline tonic signals of which 24.6% were associated with a movement within 0.2s of the onset of the change. These tonic fluctuations were highly, but not always, correlated with movement of the animal (p<0.001; **Fig. 5a-b**). Tonic cholinergic activity was also highly correlated between axon segments in the same recording session, suggesting that the fluctuations were network-wide (p<0.001; **Fig. 5c-e**) rather than in a specific sub-population. These results argue that tonic fluctuations may reflect a global change in behavioral and brain state of the animal. This global change was also reflected in the baseline activity of the cortical network as we observed a striking, temporally-correlated change in baseline cortical and axonal activity suggesting coupling between state-level changes in cortical networks and tonic cholinergic neuromodulation (p<0.001; **Supplementary Fig. 10**).

**Figure 5.**
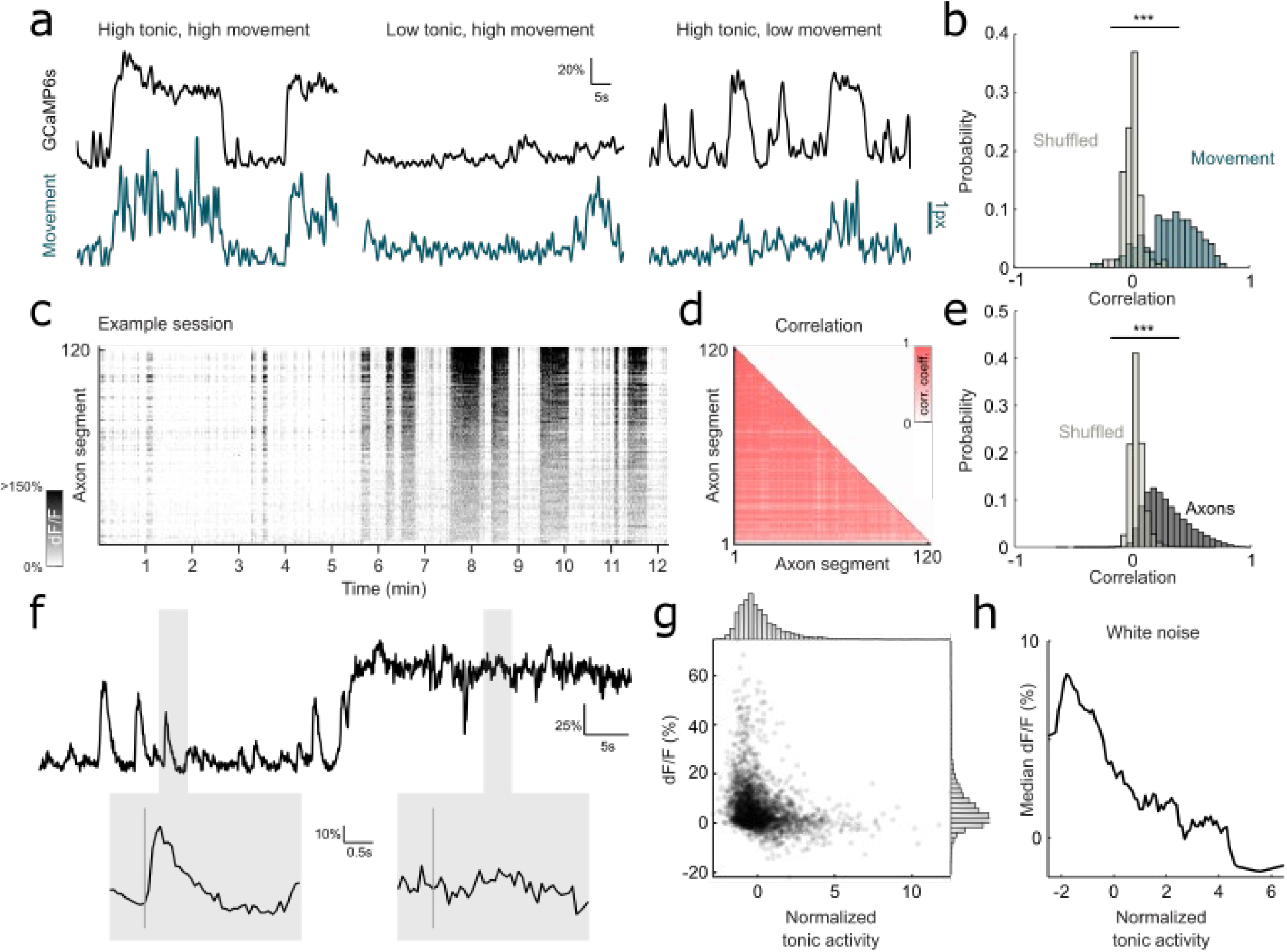
State-dependent tonic cholinergic activity modulates sound-evoked cholinergic responses. (**a**) Example tonic GCaMP6s fluorescence (black) and movement (turquoise). Some high tonic epochs are associated with movement (left), some movement are not associated with high tonic epoch (center), and some high tonic epochs are not associated with movement (right). Scalebar indicates 1-pixel movement. (**b**) Histogram of correlation coefficient of GCaMP6s signal and movement (turquoise) compared to shuffled data (gray), p<0.001. (**c**) Tonic GCaMP6s signal for all axon ROIs in example recording site. (**d**) Correlation matrix of tonic activity for all ROIs in (**c**). (**e**) Histogram of correlation coefficient of axon ROIs in each recording site (black) compared to shuffled data (gray), p<0.001. (**f**) Top: example mean fluorescence activity of one recording session showing low and high tonic activity. Shaded regions indicate response windows to white noise stimulus. Bottom: evoked response to white noise at low and high tonic activity corresponding to windows highlighted above. Gray line indicates presentation of white noise. (**g**) Scatterplot of mean evoked response to white noise at different tonic cholinergic baseline. Histogram for normalized tonic activity (top) and evoked response (right). (**h**) Median evoked response to white noise across entire dynamic range of tonic activity.

We next investigated how changes in baseline activity modulated sensory-evoked cholinergic responses. We observed that at high tonic epochs, the mean amplitudes of sound-evoked responses were significantly attenuated (**Fig. 5f**). Importantly, tonic cholinergic activity was not binary; instead, we observed a continuum of baseline activity. When we compared evoked responses to the white noise stimulus across this range of baseline cholinergic levels, we found that the amplitude of phasic cholinergic responses increased as tonic cholinergic activity ramped up to an optimal ‘sweet-spot’ and any further increase in tonic cholinergic activity led to a decrease in sound-evoked responses (**Fig. 5G-H**). Similar modulatory effects of tonic cholinergic activity were observed for pure tones and up- and down-sweeps stimuli (**Supplementary Fig. 11**). These results suggest that network-wide tonic changes in cholinergic activity (which are linked to brain and behavioral states) strongly modulates stimulus-specific sensory information relayed by phasic cholinergic signals.

## Discussion

We systematically characterized sensory-evoked responses of CBF projections to the auditory cortex. Using two-photon imaging of cholinergic axonal projections, we observed robust and non-habituating responses to auditory stimuli widely across the auditory cortex. Cholinergic sensory responses were not homogeneous, as individual axon segments displayed heterogeneous but stable tuning to pure tones. This heterogeneity allowed us to decode stimulus identity from axonal activity at a population level. Despite the response heterogeneity, cholinergic axon responses were not tonotopically organized and were largely uncoupled from the tuning of nearby cortical neurons. Chemogenetic suppression also revealed that the auditory thalamus is a primary source of auditory information from the ascending auditory pathway although this could be supplemented by inputs from earlier auditory regions (e.g. inferior colliculus or auditory brainstem). Lastly, we observed that endogenous changes in tonic cholinergic activity, reflecting both behavioral and brain states, modulates phasic sensory signaling of the CBF.

Our study demonstrates that sound-evoked cholinergic transients (1) are stably driven by repeated presentation of sounds and not merely associated with novelty or movement, (2) are intrinsically present even in the absence of behavioral conditioning, (3) encode readily the identity of the stimulus. These features argue that the CBF provides a parallel sensory channel to the auditory cortex. Interestingly, despite the heterogeneity and stimulus-specific encoding, cholinergic innervation is not tonotopically-organized and is uncoupled from cortical neural tuning. This spatial decorrelation of the parallel cholinergic sensory signal and canonical feedforward auditory signal could help calibrate cortical responses and provide a powerful substrate for experience-dependent cortical plasticity. Previous studies have shown that pairing external stimulation of basal forebrain cholinergic neurons with pure tones can induce long-lasting shifts in frequency tuning of cortical neurons^16–19^, a process achieved through the disinhibition of microcircuits by acetylcholine^18,47^. Our demonstration that cholinergic projections to the auditory cortex display intrinsic sensory responses that overlap temporally with cortical neuronal responses may provide an ecologically plausible mechanism for cortical plasticity based on sensory information from the environment. Notably, the decorrelation in tuning provides repeated representations of a broad range of sound stimuli at all points on the cortical tonotopic map, allowing cortical neurons to receive cholinergic inputs at frequencies outside of their best frequencies. This parallel channel could enable shifts in cortical tuning to behaviorally relevant stimuli which may be particularly powerful at the shoulders of a neuron’s tuning curve.

Our work also calls into question the classic dichotomy between phasic and tonic modes of neuromodulation^22,23^. The cognitive role of acetylcholine has traditionally been considered from a slow, spatially diffuse perspective based on a canonical volume transmission. Recent studies using modern experimental techniques, however, have revealed that cholinergic activity operates at multiple timescales with a more region-specific functional architecture^6,25,27,32^. Our results argue that different timescales of cholinergic activity interact in the CBF – slow cholinergic signals which indicates brain and behavioral states have profound effects on fast sensory-evoked cholinergic transients. The interaction between different modes of cholinergic signaling potentially follows a classical Yerkes-Dodson inverted-U relationship^29,48^ in which phasic sensory signals are attenuated when tonic baseline cholinergic level is too low or high, such as when the animal is overly aroused, locomoting, or disengaged. Taken together, our results suggest that the CBF is a self-regulating multiplexer, receiving sensory or task-relevant information, modulating it based on the state of the animal, and sending an integrated combination of fast and slow signal to downstream regions. Our findings serve to expand current theoretical models on the role of CBF in learning, task engagement, and decision-making and lay the groundwork for future investigation of the behavioral relevance of sensory cholinergic neuromodulation.

## Acknowledgements

We would like to thank CC, RCF, CH, DL, and SPM for helpful comments on the manuscript and CD and ZZ for assistance with cortical tonotopic measurements. This work was supported by grants from the NIH R01 DC018650, R00DC015014, NSF CAREER 2145247 and BBRF NARSAD to KVK and a JHU Science of Learning Institute Fellowship to FZ.

## Author contributions

KVK and FZ designed the study. FZ and SE performed experiments. FZ analyzed the data. JL provided analytical and conceptual advice. KVK and FZ wrote the manuscript.

## Methods

### Animals

All procedures were approved by Johns Hopkins University Animal Care and Use Committee. Male and female transgenic mice (ChAT-cre, ChAT-cre/jRGECO1a) between 6-16 weeks were used for the experiments. All experiments (passive recording and chemogenetic suppression) used ChAT-cre mice unless stated otherwise. ChAT-cre mice were obtained from The Jackson Laboratory (Stock No.: 006410) and bred in-house. ChAT-cre/jRGECO1a mice were bred in-house by crossing homozygous female ChAT-cre mice and hemizygous male jRGECO1a obtained from The Jackson Laboratory (Stock No.: 030526). First generation offspring were heterozygous for ChAT-cre and hemizygous for jRGECO1a and subsequent generation offspring were homozygous for ChAT-cre and hemizygous for jRGECO1a. Offspring genotypes were confirmed by PCR (Lucigen EconoTaq Plus GREEN 2X) and both heterozygous and homozygous ChAT-cre/jRGECO1a mice were used in the experiments and no phenotypic difference were observed.

### Surgical procedures

Mice were anesthetized with isoflurane (5.0% at induction, 2.0% during surgery) and their body temperature was maintained at 35°C throughout the surgery. For all surgeries, a 3mm craniotomy was performed over the temporal lobe (centered 1.75mm anterior to the lambda structure on the ridge line) to expose the auditory cortex. In a subset of ChAT-cre animals (n = 4) that do not endogenously express jRGECO1a in cortical neurons, an adeno-associated virus (AAV) vector encoding the calcium indicator jRGECO1a^49^ (~0.8-1.5μL, AAV1-syn-jRGECO1a, addgene) was injected in layer 2/3 in the left A1 to express calcium indicator in auditory cortical neurons. Expression of viral jRGECO1a was confirmed with two-photon microscopy. A 3mm circular glass window (Warner Instruments) was secured in place over the exposed brain with a dental cement and Krazy Glue mixture. For all animals, we carefully leveled the head of the animal and drilled a small burr hole above the basal forebrain (AP: −0.5 mm; ML: 1.8 mm; DV: 4.5 mm from bregma) and an AAV vector encoding the calcium indicator axon-GCaMP6s (1μL, AAV5-syn-flex-axon-GCaMP6s, addgene) was injected into the basal forebrain to express GCaMP6s in cholinergic neurons and their axonal projections. In animals used for chemogenetic suppression experiments, an inhibitory DREADDs hM4Di packaged into an AAV (0.8μL, AAV5-CaMKII-hM4D(Gi)-mCherry, addgene) was injected into the left medial geniculate body (n = 4; AP: −3.2mm; ML: 1.9mm; DV: −3.5mm), or left auditory cortex respectively (n = 5; 1.75mm anterior to the lambda structure on the ridge line). All injections were done using a Hamilton needle (Hamilton Company, 34 gauge, 1 inch, 12 degree bevel) and syringes (Hamilton Company, 1700 series, 5μL capacity), and a microinjection pump (Harvard Apparatus) at a flow rate of 0.60-0.75μL/min. For injections in the basal forebrain, the injection needle was left in place for at least 5 minutes following infusion to reduce backflow. Finally, a custom-made stainless steel headpost was affixed to the exposed skull with C&B Metabond dental cement (Parkell) and animals were allowed to recover for at least 3 weeks before imaging.

### Data acquisition using two-photon microscopy

Imaging was performed using a two-photon resonant-scanning microscope (Neurolabware) equipped with a 16X objective (Nikon). To image in the auditory cortex, the objective was titled to an angle of 50-60° such that it is perpendicular to the brain surface. Two-photon fluorescence of axon-GCaMP6s and jRGECO1a was excited at 980 nm using an Insight X3 laser (SpectraPhysics). We also used an electronically tunable lens to record near-simultaneously in L1 (60-100μm below dura) and L2/3 (150-200μm below dura) in sites that contained axonal segments (312μm x 192μm area, frame rate 31.92Hz overall, 15.96 per plane, laser power ≤ 40mW). As we did not observe significant differences in sound-evoked axonal response between the two layers, data across the two layers were grouped together for analysis.

To record time courses of sound-evoked axonal activity, awake animals were head-fixed under the microscope and a speaker was placed adjacent to the animal (microphone-to-ear distance ~5cm). Animals were presented with a set of 11 auditory stimuli consisting of 8 pure tones (70 dB, 4.8–54.8 kHz, half-octave intervals, 100ms, 10ms cosine on/off ramps) and 3 complex sounds (70-80 dB, white noise, frequency-modulated up-, and down-sweep, 100ms). Auditory stimuli in the set were presented in a pseudo-random order with 3.3s interval between sounds and the stimuli set was repeated 20 times during each imaging session. Scanner noise was attenuated to 40-50 dB using a custom-made foam sound enclosure directly surrounding the animal. Images were collected at 2x and 4x magnification using ScanBox software (Neurolabware) and motion-corrected with Suite2p^50^. A widefield vasculature image was also be taken at each imaging site to help with multiple site alignment.

### Data analysis

Data analysis was performed using custom functions written in MATLAB (MathWorks). To obtain time-courses of axonal and neuronal activity, we manually identified regions-of-interest (ROIs) with ImageJ (NIH) for axons and cells from mean fluorescence images at each field-of-view and extracted the timeseries of their fluorescence activity. For each presentation of auditory stimuli, we calculated ΔF/F of the sound-evoked response as the ratio of mean fluorescence in duration-matched response windows before and after tone presentation. ROIs were determined to be responsive to a particular stimulus if their evoked responses showed a significant difference across 20 presentations of the same stimuli (p<0.025, right-tailed paired t-test).

To align multiple sites in each animal, pixel-wise x- and y-offset between each imaging site were measured by manually comparing vasculature images using Photoshop v14.0 (Adobe). These offset values were used in a custom MATLAB function to stitch the vasculature and two-photon images together. For analysis of axonal tonotopy in the primary auditory cortex, the primary auditory cortex was first located by analyzing cortical neuronal (jRGECO1a) response for imaging sites with tone-responsive neurons. The relative positions of axon segments in the primary auditory cortex along the rostro-caudal axis were obtained from the stitched image and plotted against their most responsive frequency. Tonotopy is operationalized as the change in best frequency of cholinergic axon segments along the rostro-caudal axis. To compare tonotopy between cholinergic axons and cortical neurons, size-matched area of primary auditory cortex were identified in animals expressing the same family of calcium indicator (GCaMP6f) in excitatory cortical neurons. These animals underwent the same surgical process described above but received viral injection of GCaMP6f (1μL, AAV9-CamKII-GCaMP6f, addgene) in the same coordinates in the auditory cortex and did not receive axon-GCaMP6s injection in the basal forebrain. The primary auditory cortex was located in these animals by identifying the region with an increasing change in best frequency along the rostro-caudal axis as described in previous studies^22^. Tonotopy of cortical neurons were quantified as described above.

For comparison of cortical neuron and axonal tuning, distance of each ROI was calculated as the Euclidian distance between the center of the ROIs. ROIs within 20μm were considered as ‘nearby’. As we were unable to accurately determine the z-offset between each imaging site, cortical neurons and nearby axonal segments and neurons used were limited to within each imaging site. To improve signal-to-noise ratio for analysis comparing tuning of cortical neurons and nearby cortical neurons, analysis was restricted to cell ROIs with evoked response greater than the noise ceiling (97.5^th^ percentile of all fluorescence activity).

For tonic activity correlation analysis, a lowpass filter (passband frequency = 0.5Hz) was applied to the raw fluorescence trace and the movement signal. Correlation coefficient is calculated for the relevant filtered timeseries using the entire session. Movement was calculated using the x-y offset of the motion-corrected image. x-y offset was extracted using Suite2p and the amplitude of movement signal was calculated as the absolute difference of the Euclidean norm of x- and y-offset for each successive frame. To quantify tonic fluctuations that were closely coupled with movement, changes in tonic activity and movement were digitized using respective thresholds. The tonic threshold was defined as two median absolute deviations above median tonic activity of each recording session; the movement threshold was defined as x-y offset greater than 1 pixel. Tonic epochs were labeled as closely coupled with movement if onset of movement occur within 0.2s of change in tonic activity. Processed data were visually inspected to validate the appropriateness of the chosen thresholds. To compare tonic cholinergic activity across imaging sessions and animals, fluorescence of each session was standardized by subtracting the median and dividing this difference by the median absolute deviation. This method of standardization was adopted as we observed a wide dynamic range of baseline tonic activity that could not be digitally classified into ‘low’ and ‘high’. On this interval scale, median level of tonic activity is designated ‘0’, whereas low tonic epochs are negative and high tonic epochs are positive. This allowed us to compare tonic cholinergic activity without setting an arbitrary ‘tonic floor’.

For multi-class decoding, we used a naïve Bayes classifier to classify calcium activity into multiple stimuli classes. We trained the frame-by-frame decoder using frame-by-frame raw fluorescence values of all axon ROIs for 19 presentations of the three complex auditory stimuli or eight pure tone and tested the decoder on a left-out trial. We validated stimulus-decoding accuracy with a twenty-fold cross-validation. Shuffled data was constructed from the same axonal activity but the label for tone identity was randomized. 95% confidence interval for shuffled data was calculated by iterating the classification of shuffled data for 100 times and taking the value of the 2.5^th^ and 97.5^th^ percentile. To investigate if performance of the linear decoder was driven by high decoding accuracy of specific tones, we conducted pairwise decoding using the same naïve Bayes classifier applied to every pair of auditory stimuli (complex sounds or pure tones). We trained the decoder with mean raw fluorescence values of the frames with maximum decoding as determined by the previous analysis (3-7 frames after tone presentation) of all axon segments. To test the robustness of our decoding, we trained our decoders with population activity from all axon ROIs and tested their decoding accuracy while removing the top n^th^ percentile of most influential ROIs (based on the size of the weights). We further examined decoding accuracy per animal by training the frame-by-frame and pairwise decoder on responsive axon ROI activity in 6 animals with more than 100 responsive axon segments.

### Chemogenetic suppression

Mice expressing inhibitory DREADDs hM4Di first received 10mL/kg intraperitoneal injections of saline. 15min after saline injection, the animals were placed under the two-photon microscope and activity of cholinergic axonal projections to the auditory cortex was recorded in a similar protocol described above. At the end of the imaging session, animals were removed from head-fixation for 5min before receiving intraperitoneal injection of 0.5-3mg/kg clozapine N-oxide (CNO). Volume of saline and CNO injections were matched. 15min after CNO injection, the animals were placed back under the two-photon microscope and activity of cholinergic axonal projections to the auditory cortex was again recorded. Efforts were made to image the same axons for saline and CNO injections. At the end of the experiment, a subset of mice was perfused for histology to determine the expression of hM4Di. Recording sessions for saline and CNO injections were aligned and preprocessed separately and the responses of cholinergic axon segments were quantified as described above. Main effect of CNO injection was quantified using 2-way ANOVA (Type II SS). Analyses comparing mean evoked response after saline and CNO injection were limited to 9.5-19kHz as these tones elicited evoked responses in the cholinergic axons in the imaging sites following saline injection.

To verify that CNO injection suppressed the medial geniculate body and auditory cortex in mice expressing hM4Di in the respective areas, control experiments were conducted. ChAT-cre mice received GCaMP6f injection in the auditory cortex (1μL, AAV9-CamKII-GCaMP6f, addgene) and hM4Di injection in either the medial geniculate body or auditory cortex as described above. 3 weeks after injections, chemogenetic suppression protocol described above were conducted and cortical response to auditory stimuli were recorded following intraperitoneal saline and CNO injection. Preprocessing and quantification of cortical responses were performed as described above. Analyses comparing mean evoked response after saline and CNO injection were limited to 9.5-19kHz for medial geniculate body suppression condition and 4.8-19kHz auditory cortex suppression condition as these tones elicited evoked responses in the cortical neurons in the imaging sites following saline injection.

### Histology

To confirm the specific expression of axon-GCaMP6s in basal forebrain cholinergic neurons following injection in ChAT-cre mice, we performed immunohistochemistry with ChAT and GFP antibodies. We also performed histological analysis (without antibodies) to confirm the expression of inhibitory DREADDs hM4Di (which expresses a mCherry fluorescence marker) in neurons in the medial geniculate body and auditory cortex respectively.

Mice were deeply anesthetized and transcardially perfused with ~20mL phosphate-buffered saline (PBS) solution followed by ~20mL 4% PFA. Brains were then extracted from the skull and post-fixed in 4% PFA overnight at 4°C before transfer to 30% sucrose solution for 2-3 days at 4°C. Next, the brains were frozen in tissue tek O.C.T. compound (Sakura Finetek) at 80°C for multiple days to prepare for slicing. Frozen brains were sliced coronally with 35μm thickness on a cryostat and permeabilized for 15min with 0.3%PBS-Triton (PBS solution with 0.3% Triton X-100 (Sigma Aldrich)). Slices were incubated for 1hr in a blocking buffer containing 0.3% PBS-Triton and 10% Normal Donkey Serum (Synaptic Systems). Slices were then transferred to fresh 0.3% PBS-Triton and incubated overnight at 4°C with appropriate primary antibodies (1:200-500 dilution of goat anti-ChAT IgG, Millipore, AB114P; 1:500 rabbit anti-GFP IgG, Abcam, ab6556 or 1:300 rabbit anti-GFP IgG, ThermoFisher, A-6455 (both anti-GFP had similar level of expression)).

Afterwards, slices were washed in PBS solution and incubated for 1hr at room temperature with secondary antibody (1:500 Cy™3 AffiniPure Donkey Anti-Goat IgG, Jackson ImmunoResearch, 705-165-147; 1:500 Alexa Fluor^®^ 488 AffiniPure Donkey Anti-Rabbit IgG, Jackson ImmunoResearch, 711-545-152). Finally, slices were rinsed in PBS solution and incubated at room temperature in DAPI Fluoromount-G (Southern Biotech) before being mounted onto glass slides and coverslipped for imaging.

Images for cell counting were acquired using a 20x air objective on a Zeiss LSM 700 Confocal Microscope (Carl Zeiss) from the basal forebrain for axon-GCaMP6s immunohistochemistry. Cell counts were performed manually in ImageJ (NIH). Coronal slice images were acquired using a 10x air objective on a Zeiss LSM 700 Confocal Microscope (Carl Zeiss). The basal forebrain, medial geniculate nucleus and auditory cortex were located using coordinates from the Allen Brain Atlas and references from other studies^41,42^.

### Statistical Analysis

All statistical analyses were performed in MATLAB (MathWorks). All data are reported as mean ± SEM unless otherwise indicated. Statistical significance was defined as p<0.05 unless otherwise indicated.

## Supplementary Information

**Supplementary Fig. 1.**
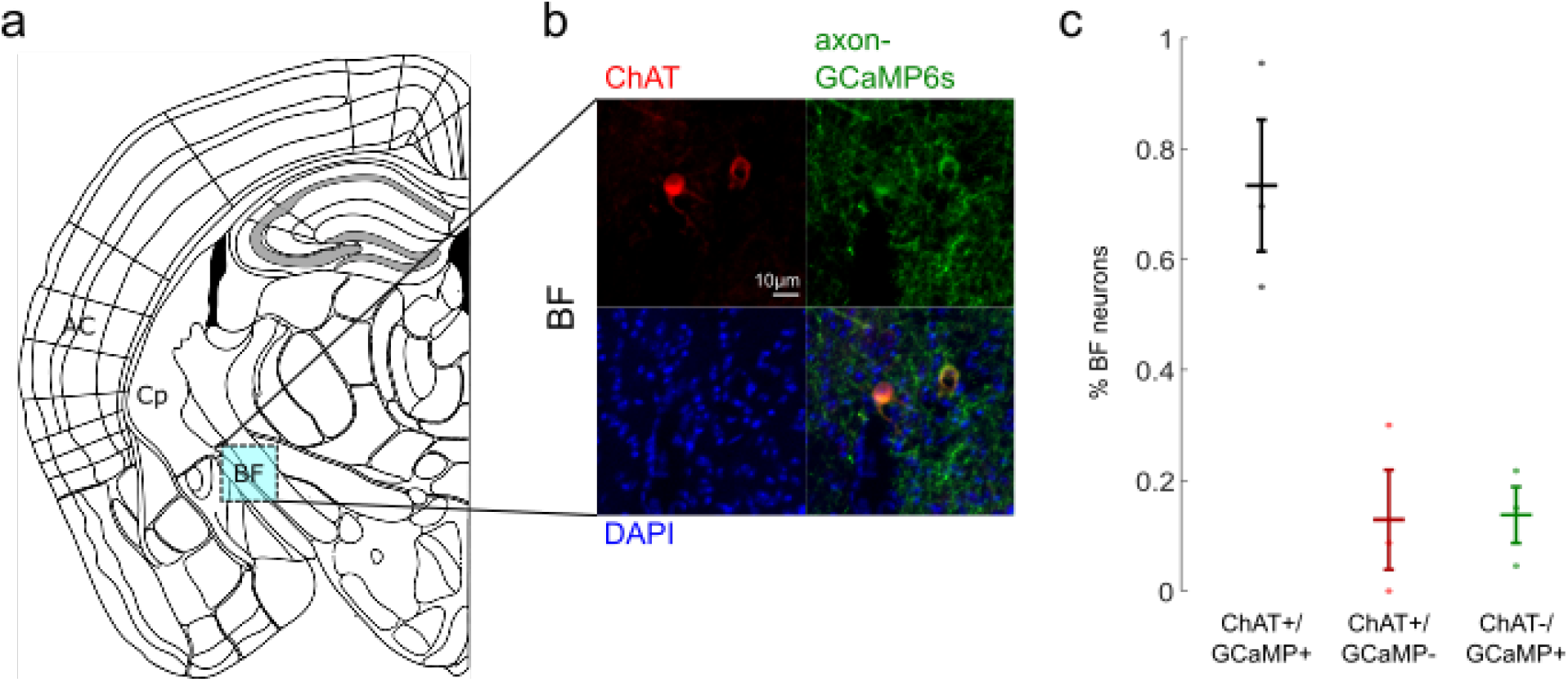
Histology for cre-dependent cholinergic neurons targeting. (**a**) Schematic of imaging site for basal forebrain (BF). (**b**) Basal forebrain stained for inhibitory ChAT (red), axon-GCaMP6s (green), and DAPI. Histology is validated in 3 animals. (**c**) Percentage of basal forebrain neurons that express both axon-GCaMP6s and ChAT (black), ChAT-only (red), or axon-GCaMP6s-only (green).

**Supplementary Fig. 2.**
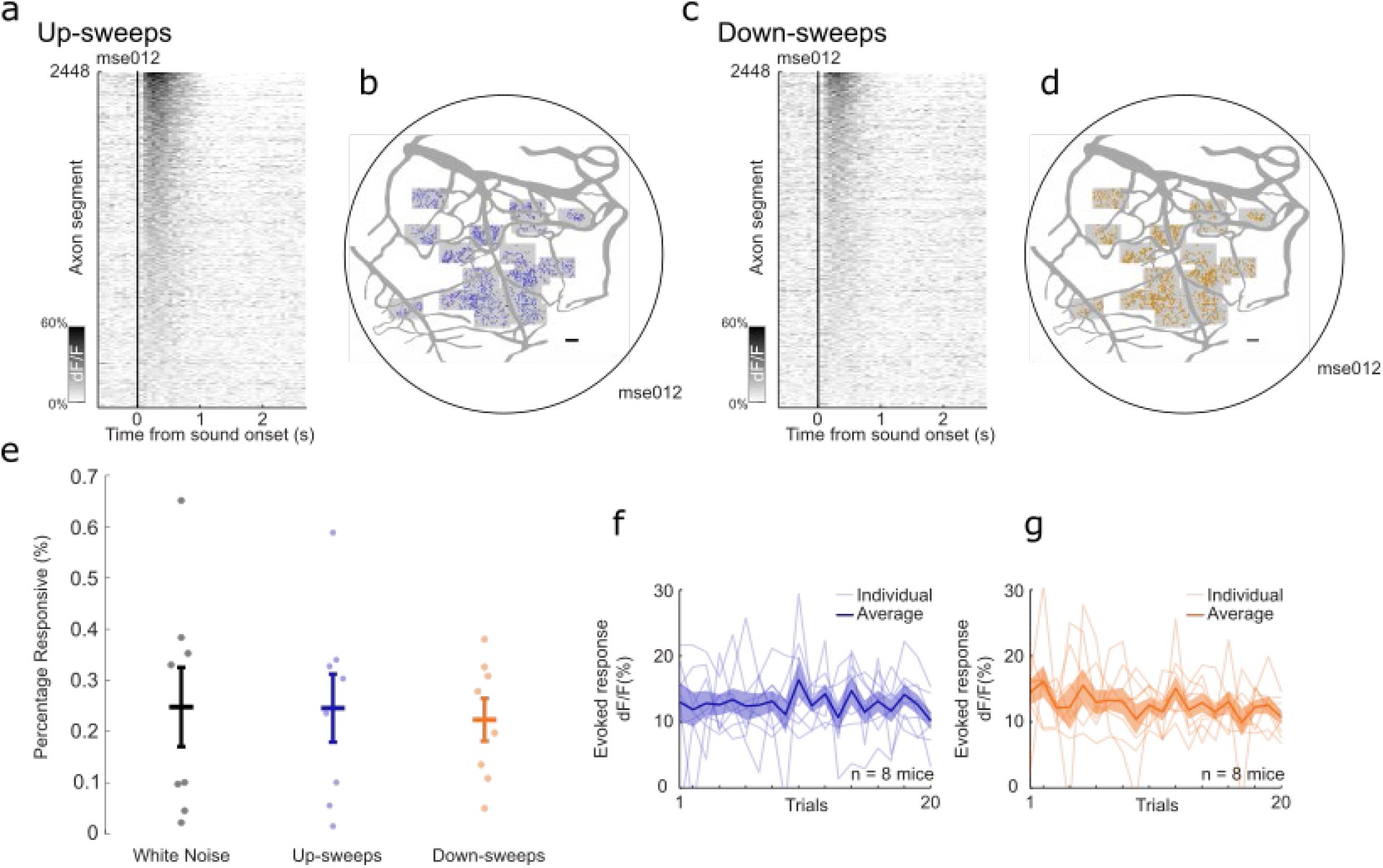
Robust and non-habituating response to up-sweeps and downs-weeps. (**a**) Heatmap of average evoked response (ΔF/F) to up-sweeps for all identified axon segments in one animal (n = 2448 axon segments). (**b**) Spatial distribution of axon segments responsive to up-sweeps (blue) in one animal. Shaded boxes indicate recording sites. Scalebar = 100μm (**c**) Heatmap of average evoked response (ΔF/F) to down-sweeps for all identified axon segments in one animal (n = 2448 axon segments). (**d**) Spatial distribution of axon segments responsive to down-sweeps (orange) in one animal. Shaded boxes indicate recording sites. Scalebar = 100μm (**e**) Percentage of identified axon segments that are responsive to white noise (black), up-sweeps (blue), and down-sweeps (orange) in 8 animals (**f**) Amplitude of evoked response for up-sweeps across 20 presentations for all animals (n = 8 animals). (**g**) Amplitude of evoked response for down-sweeps across 20 presentations for all animals (n = 8 animals).

**Supplementary Fig. 3.**
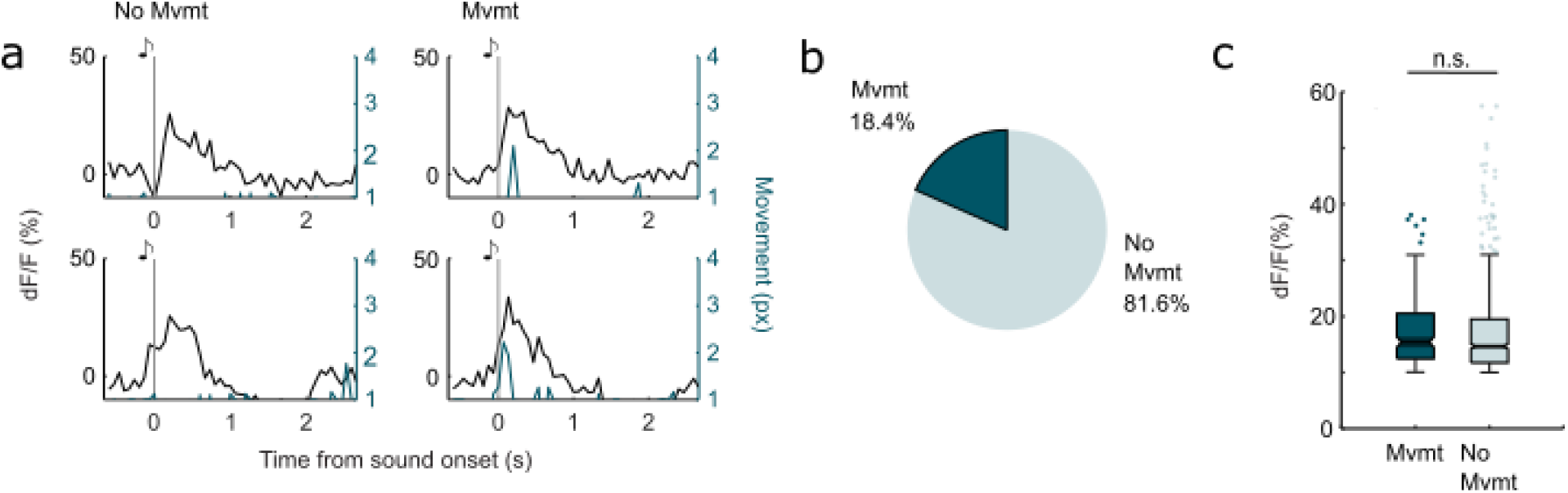
Micromovements are associated with some but not all phasic cholinergic transients. (**a**) Example stimulus-synchronous phasic cholinergic transients from one example axon ROI that are associated with micromovement (left) adtuned to nearby cortical neuronsn not associated with micromovement (right). (**b**) 18.4% of stimulus-synchronous phasic transients are associated with micromovements. (**c**) Micromovement does not significantly modulate amplitude of sound-evoked transients, p = 0.554.

**Supplementary Fig. 4.**
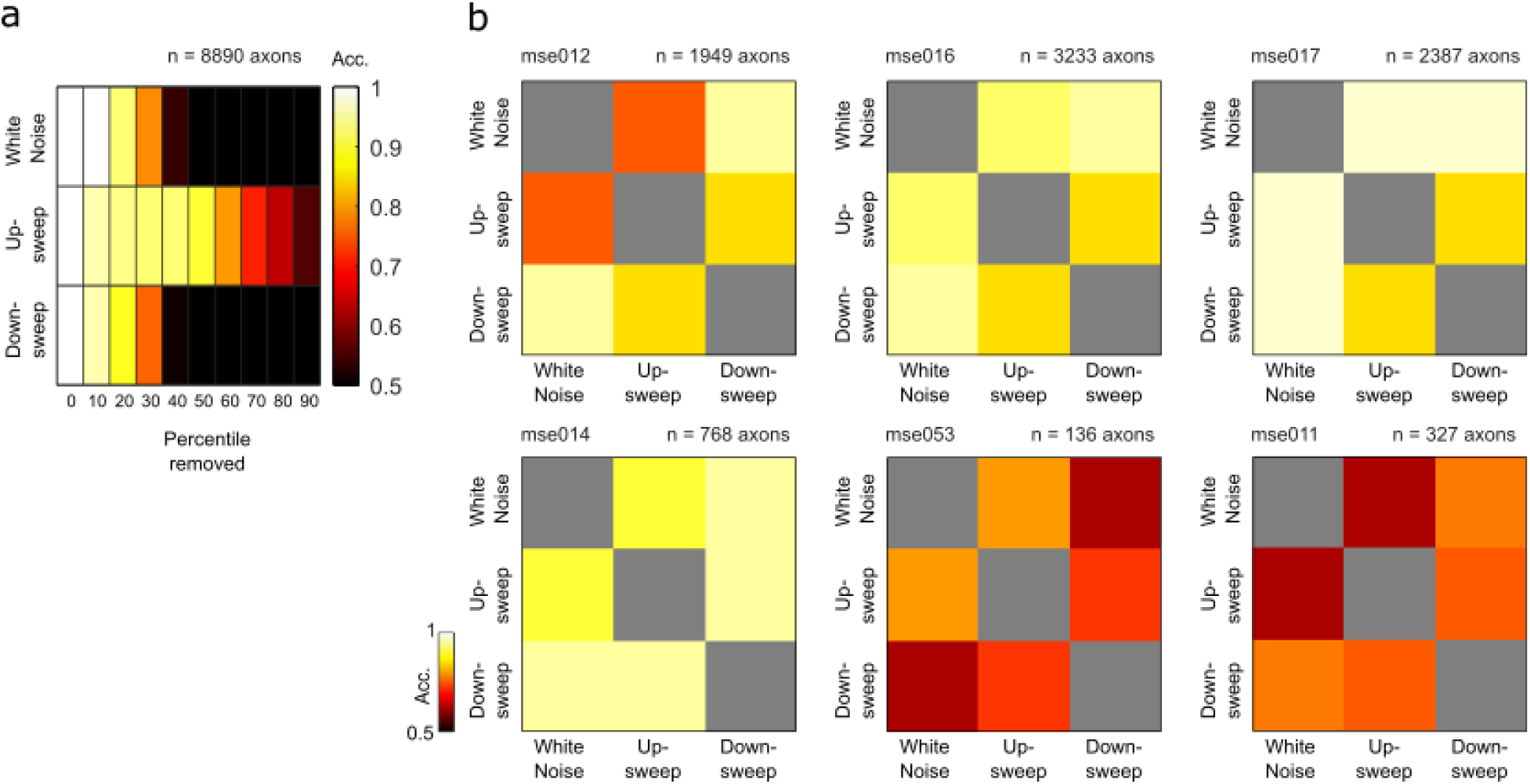
Robust stimulus-specific decoding of complex sounds. (**a**) Average pairwise decoding accuracy for each complex sound stimulus removing n^th^ percentile of most influential ROIs. (**b**) Pairwise decoder accuracy for complex sound stimuli on population activity of responsive axon segments in animals with more than 100 responsive axon segments. All sound-pairs are significantly above chance.

**Supplementary Fig. 5.**
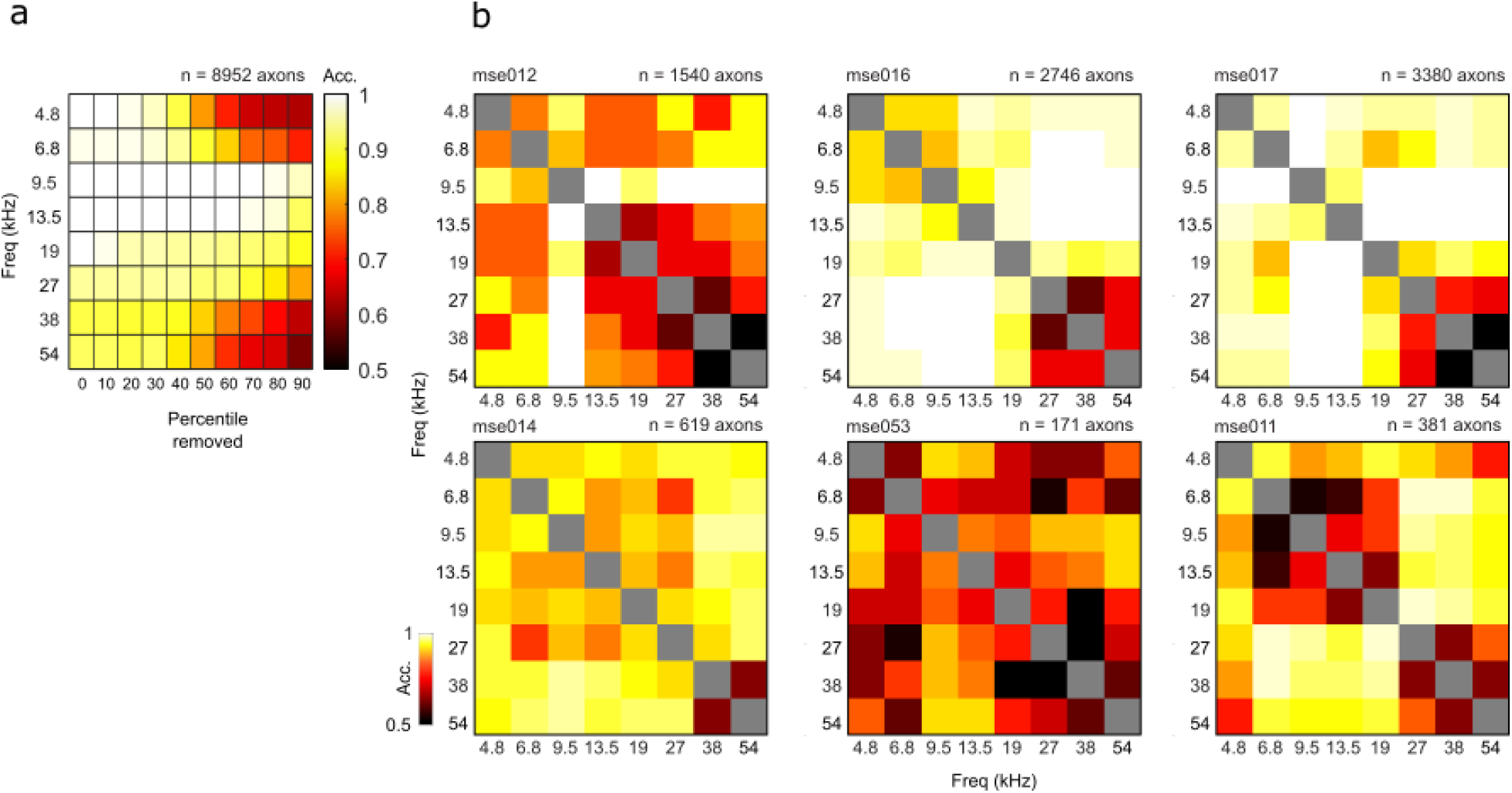
Robust stimulus-specific decoding of pure tones. (**a**) Average pairwise decoding accuracy for each pure tone removing n^th^ percentile of most influential ROIs. (**b**) Pairwise decoder accuracy for pure tones on population activity of responsive axon segments in animals with more than 100 responsive axon segments. 97.6±0.1% of sound-pairs are significantly above chance.

**Supplementary Fig. 6.**
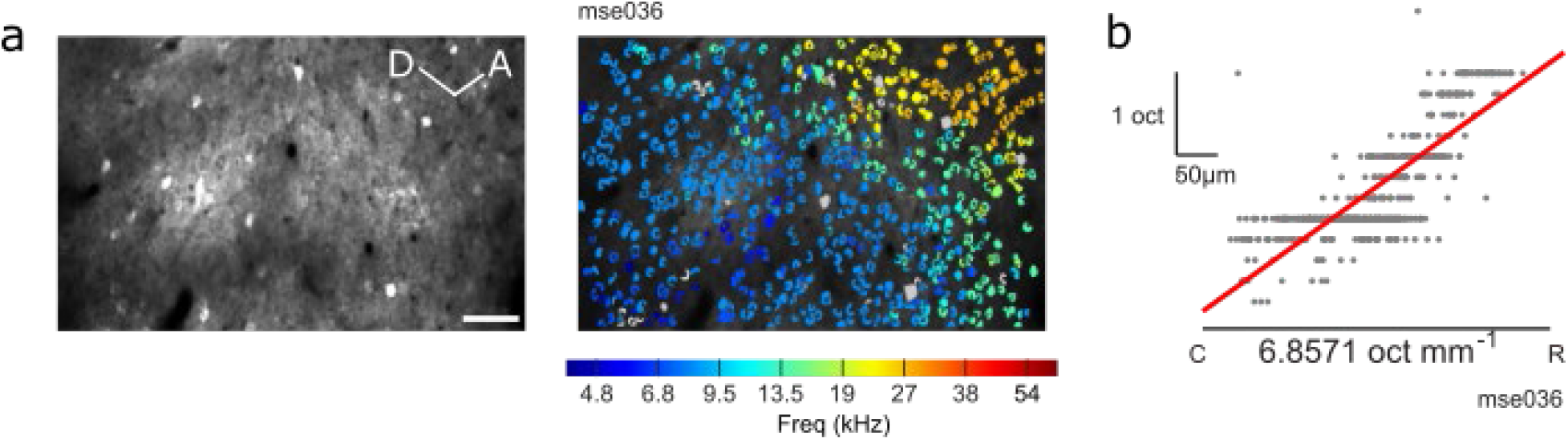
Tonotopic gradient of excitatory neurons in primary auditory cortex. (**a**) Example field-of-view of cortical neurons in primary auditory cortex (left, CaMKII-GCaMP6f) and identified ROIs colored by best frequency of cortical neurons (right). Scalebar = 50μm (**b**) Change in best frequency of neuron ROIs in (**a**) along the caudal-rostral axis.

**Supplementary Fig. 7.**
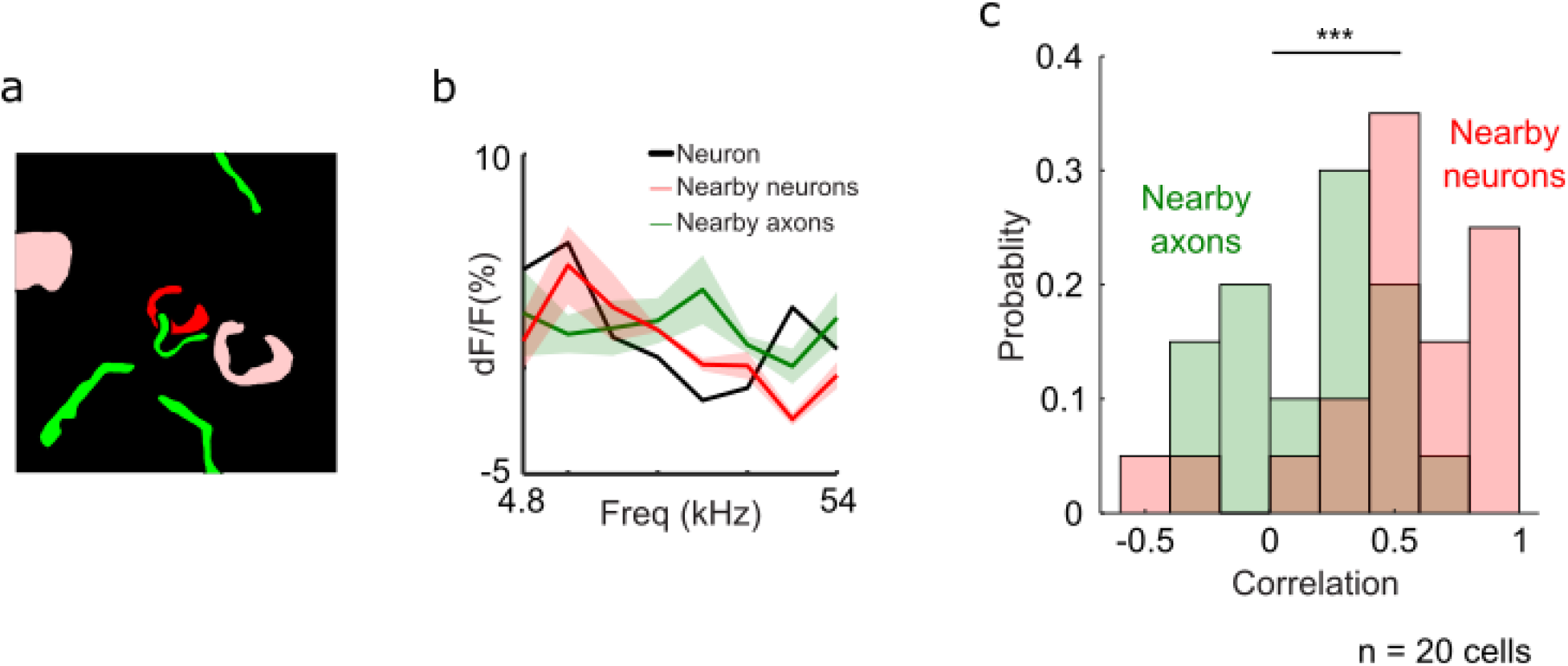
Cortical neurons are co-tuned to nearby cortical neurons but un-coupled from nearby cholinergic axons. (**a**) Schematic of example neuron (red) and nearby neurons (pink) and responsive axon segments (green) (within 20μm). (**b**) Frequency tuning curve of example neuron (black) and nearby neurons (red) and axon segments (green) in (**a**). (**c**) Histogram of correlation coefficient between tuning of auditory cortical neurons with nearby cortical neurons (red) and nearby axon segments (green).

**Supplementary Fig. 8.**
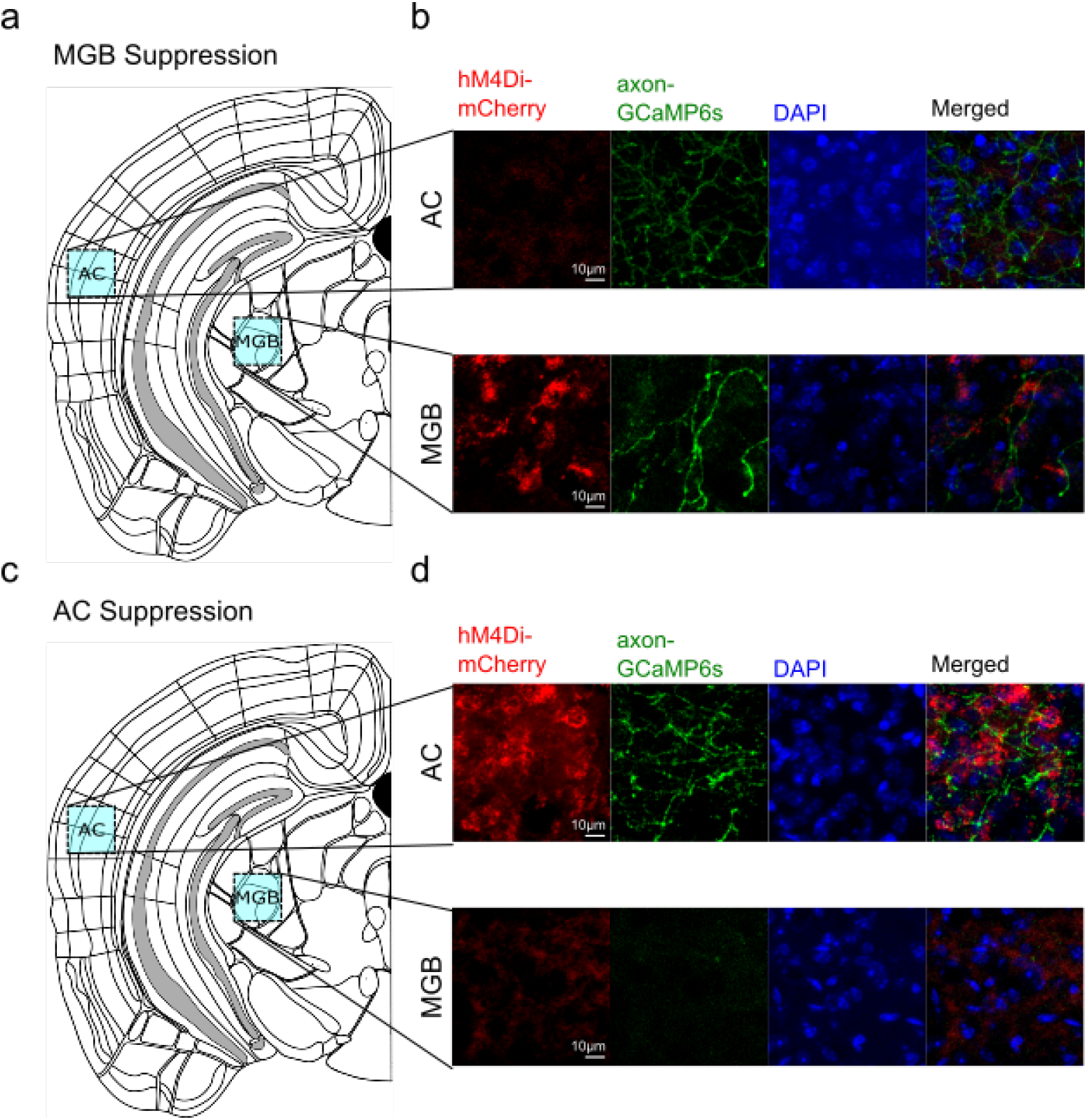
Histology for medial geniculate body and auditory cortex DREADDs targeting. (**a**) Schematic of imaging site for auditory cortex (AC) and medial geniculate body (MGB) in MGB suppression mice. (**b**) Top: auditory cortex stained for inhibitory DREADDs hM4Di (red), axon-GCaMP6s (green), and DAPI. Bottom: medial geniculate body stained for inhibitory DREADDs hM4Di (red), axon-GCaMP6s (green), and DAPI. Histology is validated in 4 experimental animals. (**c**) Schematic of imaging site for auditory cortex (AC) and medial geniculate body (MGB) in AC suppression mice. (**d**) Top: auditory cortex stained for inhibitory DREADDs hM4Di (red), axon-GCaMP6s (green), and DAPI. Bottom: medial geniculate body stained for inhibitory DREADDs hM4Di (red), axon-GCaMP6s (green), and DAPI. Histology is validated in 2 experimental animals.

**Supplementary Fig. 9.**
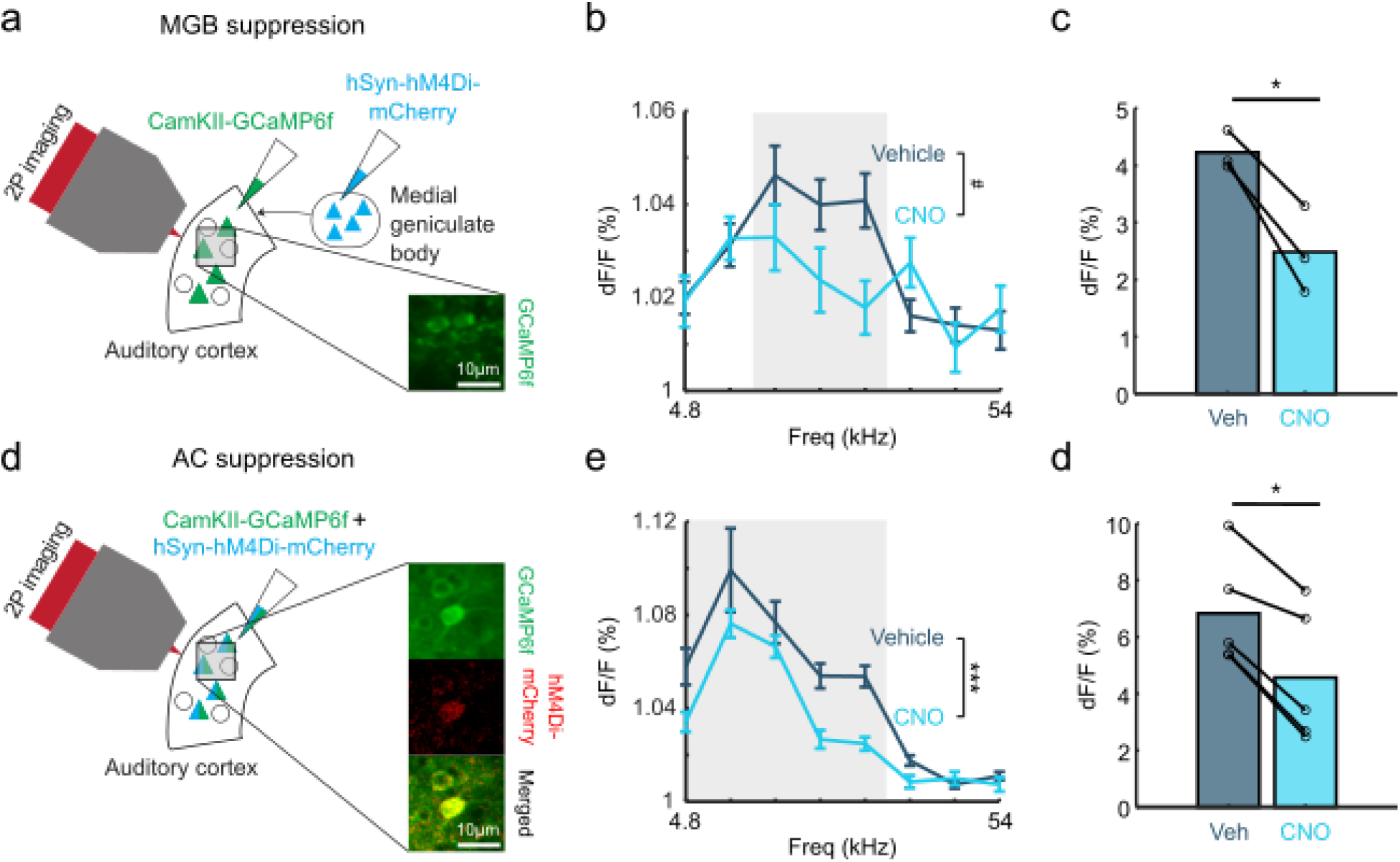
Chemogenetic suppression of auditory thalamus and auditory cortex attenuates sound-evoked cortical responses. (**a**) Schematic of injection strategy for suppression of the medial geniculate body. Inset: cortical neurons expressing GCaMP6f (green) (**b**) Evoked cortical response to pure tones after intraperitoneal saline and CNO injection (n = 95 cells for saline condition; n = 55 cells for CNO condition, F(1,1184) = 3.57, p = 0.0589). Shaded region significantly responsive tones identified post saline injection (9.5-18kHz). (**c**) Mean evoked response after intraperitoneal saline and CNO injection for each significantly responsive tone, p<0.05. (**d**) Schematic of injection strategy for suppression of the auditory cortex. Inset: cortical neurons expressing GCaMP6f (green), inhibitory DREADDs hM4Di (red) and overlaid image. (**e**) Evoked cortical response to pure tones after intraperitoneal saline and CNO injection (n = 232 cells for saline condition; n = 113 cells for CNO condition, F(1,2744) = 13.34, p<0.001). Shaded region represents significantly responsive tones identified post saline injection (4.8-19kHz). (**f**) Mean evoked response after intraperitoneal saline and CNO injection for each significantly responsive tone, p<0.05.

**Supplementary Fig. 10.**
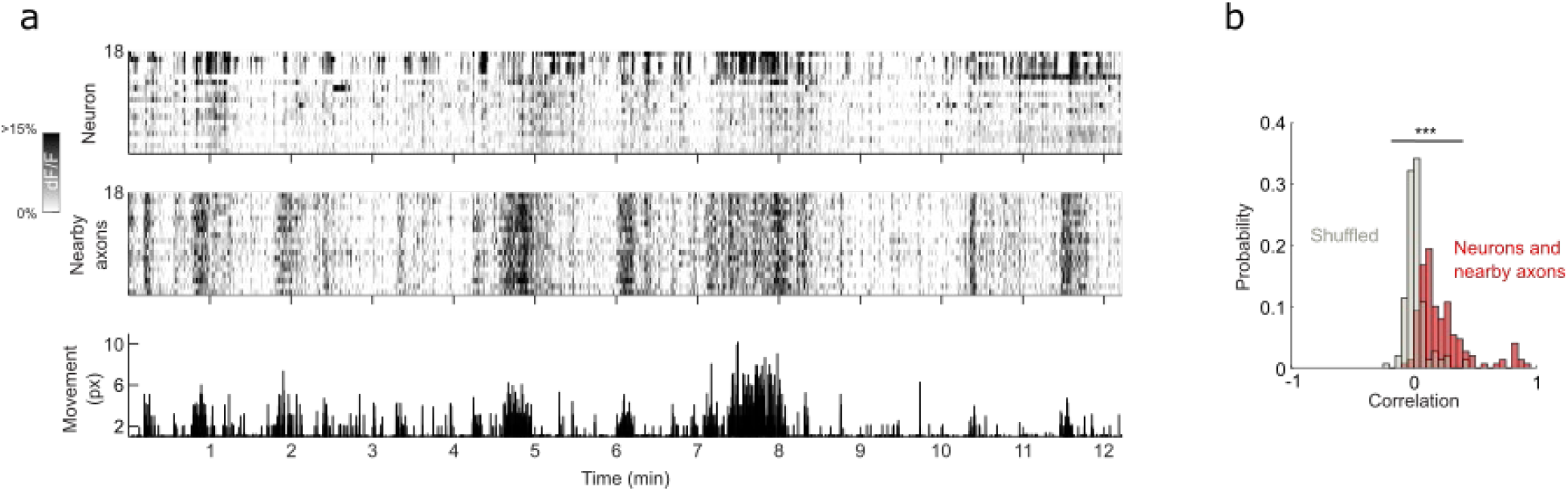
Strong coupling between local tonic cholinergic activity and tonic cortical neuron activity. (**a**) Fluorescence activity of neurons in one example recording site (top) and the nearby axons of the respective neurons (middle) and movement of the animal during the recording session (bottom). (**b**) Histogram of correlation coefficient of cell tonic activity and tonic activity of nearby axons (red) compared to shuffled data (gray), p<0.001.

**Supplementary Fig. 11.**
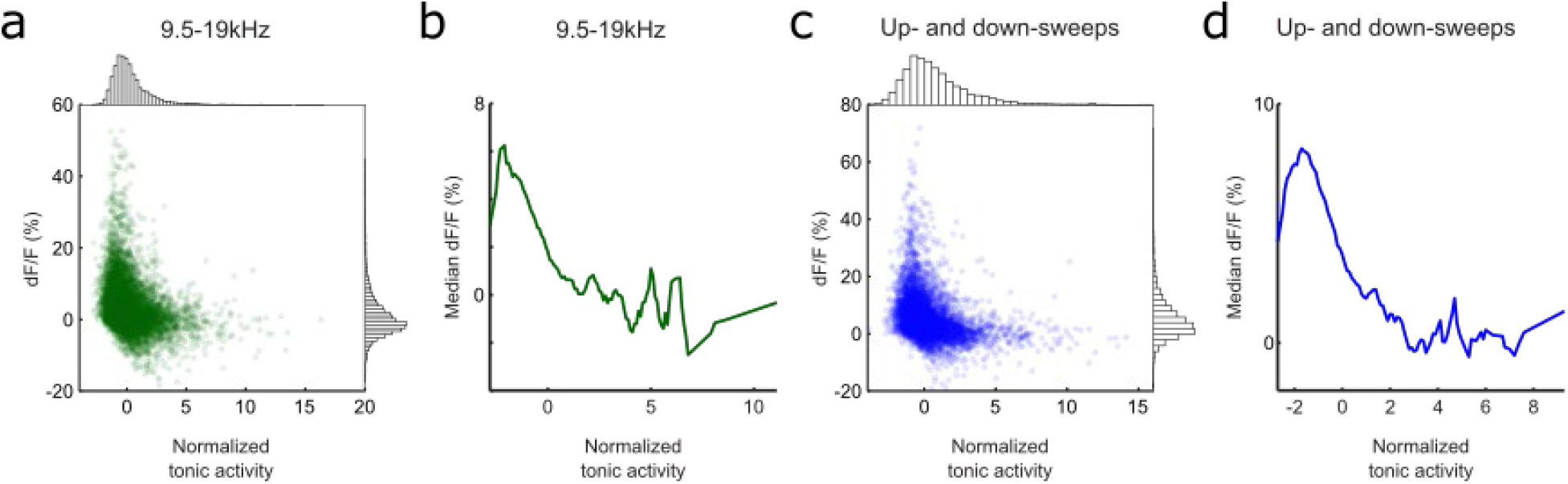
State-dependent tonic cholinergic activity modulates cholinergic response to pure tones and up- and down-sweeps. (**a**) Scatterplot of mean evoked response to 9.5-19kHz at different tonic cholinergic baseline. Histogram for normalized tonic activity (top) and evoked response (right). (**b**) Median evoked response to 9.5-19kHz across entire dynamic range of tonic activity. (**c**) Scatterplot of mean evoked response to up- and down-sweeps at different tonic cholinergic baseline. Histogram for normalized tonic activity (top) and evoked response (right). (**d**) Median evoked response to up- and down-sweeps across entire dynamic range of tonic activity.

## References

1. Mesulam, M. M., Mufson, E. J., Wainer, B. H. & Levey, A. I. Central cholinergic pathways in the rat: An overview based on an alternative nomenclature (Ch1-Ch6). Neuroscience 10, 1185–1201 (1983).

2. Everitt, B. J. & Robbins, T. W. Central Cholinergic Systems and Cognition. Annu. Rev. Psychol. 48, 649–684 (1997).

3. Zaborszky, L., Gaykema, R. P., Swanson, D. J. & Cullinan, W. E. Cortical input to the basal forebrain. Neuroscience (1997).

4. Gielow, M. R. & Zaborszky, L. The Input-Output Relationship of the Cholinergic Basal Forebrain. Cell Rep. (2017).

5. Chavez, C. & Zaborszky, L. Basal Forebrain Cholinergic-Auditory Cortical Network: Primary Versus Nonprimary Auditory Cortical Areas. Cereb. Cortex 27, 2335–2347 (2017).

6. Parikh, V., Kozak, R., Martinez, V. & Sarter, M. Prefrontal Acetylcholine Release Controls Cue Detection on Multiple Timescales. Neuron 56, 141–154 (2007).

7. Lin, S. C. & Nicolelis, M. A. L. Neuronal Ensemble Bursting in the Basal Forebrain Encodes Salience Irrespective of Valence. Neuron 59, 138–149 (2008).

8. Higley, M. J. & Picciotto, M. R. Neuromodulation by acetylcholine: Examples from schizophrenia and depression. Curr. Opin. Neurobiol. 29, 88–95 (2014).

9. Gritton, H. J. et al. Cortical cholinergic signaling controls the detection of cues. Proc. Natl. Acad. Sci. U. S. A. 113, E1089–E1097 (2016).

10. Sarter, M. & Lustig, C. Cholinergic double duty: cue detection and attentional control. Curr. Opin. Psychol. 29, 102–107 (2019).

11. Hasselmo, M. E. The role of acetylcholine in learning and memory. Current Opinion in Neurobiology (2006).

12. Newman, E. L., Gupta, K., Climer, J. R., Monaghan, C. K. & Hasselmo, M. E. Cholinergic modulation of cognitive processing: Insights drawn from computational models. Front. Behav. Neurosci. 6, 1–19 (2012).

13. Letzkus, J. J. et al. A disinhibitory microcircuit for associative fear learning in the auditory cortex. Nature 480, 331–335 (2011).

14. Ballinger, E. C., Ananth, M., Talmage, D. A. & Role, L. W. Basal Forebrain Cholinergic Circuits and Signaling in Cognition and Cognitive Decline. Neuron 91, 1199–1218 (2016).

15. Maurer, S. V. & Williams, C. L. The cholinergic system modulates memory and hippocampal plasticity via its interactions with non-neuronal cells. Front. Immunol. 8, (2017).

16. Bakin, J. S. & Weinberger, N. M. Induction of a physiological memory in the cerebral cortex by stimulation of the nucleus basalis. Proc. Natl. Acad. Sci. U. S. A. 93, 11219–11224 (1996).

17. Kilgard, M. P. & Merzenich, M. M. Cortical map reorganization enabled by nucleus basalis activity. Science. 279, 1714–1718 (1998).

18. Froemke, R. C., Merzenich, M. M. & Schreiner, C. E. A synaptic memory trace for cortical receptive field plasticity. Nature 450, 425–429 (2007).

19. Froemke, R. C. et al. Long-term modification of cortical synapses improves sensory perception. Nat. Neurosci. 16, 79–88 (2013).

20. Takesian, A. E., Bogart, L. J., Lichtman, J. W. & Hensch, T. K. Inhibitory circuit gating of auditory critical-period plasticity. Nat. Neurosci. 21, 218–227 (2018).

21. Sarter, M., Lustig, C., Howe, W. M., Gritton, H. & Berry, A. S. Deterministic functions of cortical acetylcholine. Eur. J. Neurosci. 39, 1912–1920 (2014).

22. Disney, A. A. & Higley, M. J. Diverse Spatiotemporal Scales of Cholinergic Signaling in the Neocortex. J. Neurosci. (2020).

23. Sarter, M. & Lustig, C. Forebrain Cholinergic Signaling: Wired and Phasic, Not Tonic, and Causing Behavior. J. Neurosci. 40, 712–719 (2020).

24. Do, J. P. et al. Cell type-specific long-range connections of basal forebrain circuit. Elife 5, 1–18 (2016).

25. Kim, J. H. et al. Selectivity of neuromodulatory projections from the basal forebrain and locus ceruleus to primary sensory cortices. J. Neurosci. 36, 5314–5327 (2016).

26. Yang, C., Thankachan, S., McCarley, R. W. & Brown, R. E. The menagerie of the basal forebrain: how many (neural) species are there, what do they look like, how do they behave and who talks to whom? Curr. Opin. Neurobiol. 44, 159–166 (2017).

27. Laszlovszky, T. et al. Distinct synchronization, cortical coupling and behavioral function of two basal forebrain cholinergic neuron types. Nat. Neurosci. 23, 992–1003 (2020).

28. Buzsaki, G. et al. Nucleus basalis and thalamic control of neocortical activity in the freely moving rat. J. Neurosci. (1988).

29. McGinley, M. J. et al. Waking State: Rapid Variations Modulate Neural and Behavioral Responses. Neuron 87, 1143–1161 (2015).

30. Reimer, J. et al. Pupil fluctuations track rapid changes in adrenergic and cholinergic activity in cortex. Nat. Commun. 7, 1–7 (2016).

31. Kuchibhotla, K. V. et al. Parallel processing by cortical inhibition enables context-dependent behavior. Nat. Neurosci. 20, 62–71 (2017).

32. Teles-Grilo Ruivo, L. M. et al. Coordinated Acetylcholine Release in Prefrontal Cortex and Hippocampus Is Associated with Arousal and Reward on Distinct Timescales. Cell Rep. 18, 905–917 (2017).

33. Lohani, S. et al. Dual color mesoscopic imaging reveals spatiotemporally heterogeneous coordination of cholinergic and neocortical activity. bioRxiv (2021).

34. Hangya, B., Ranade, S. P., Lorenc, M. & Kepecs, A. Central Cholinergic Neurons Are Rapidly Recruited by Reinforcement Feedback. Cell 162, 1155–1168 (2015).

35. Harrison, T. C., Pinto, L., Brock, J. R. & Dan, Y. Calcium imaging of basal forebrain activity during innate and learned behaviors. Front. Neural Circuits 10, 1–12 (2016).

36. Crouse, R. B. et al. Acetylcholine is released in the basolateral amygdala in response to predictors of reward and enhances the learning of cue-reward contingency. Elife 9, 1–31 (2020).

37. Sturgill, J. F. et al. Basal forebrain-derived acetylcholine encodes valence-free reinforcement prediction error. bioRxiv (2020).

38. Pinto, L. et al. Fast modulation of visual perception by basal forebrain cholinergic neurons. Nat. Neurosci. (2013).

39. Eggermann, E., Kremer, Y., Crochet, S. & Petersen, C. C. H. Cholinergic Signals in Mouse Barrel Cortex during Active Whisker Sensing. Cell Rep. (2014).

40. Nelson, A. & Mooney, R. The Basal Forebrain and Motor Cortex Provide Convergent yet Distinct Movement-Related Inputs to the Auditory Cortex. Neuron 90, 635–648 (2016).

41. Guo, W., Robert, B. & Polley, D. B. The Cholinergic Basal Forebrain Links Auditory Stimuli with Delayed Reinforcement to Support Learning. Neuron 103, 1164–1177.e6 (2019).

42. Robert, B. et al. A functional topography within the cholinergic basal forebrain for encoding sensory cues and behavioral reinforcement outcomes. Elife 10, 1–28 (2021).

43. Zhang, K., Chen, C. D. & Monosov, I. E. Novelty, Salience, and Surprise Timing Are Signaled by Neurons in the Basal Forebrain. Curr. Biol. 29, 134–142.e3 (2019).

44. Stiebler, I., Neulist, R., Fichtel, I. & Ehret, G. The auditory cortex of the house mouse: Left-right differences, tonotopic organization and quantitative analysis of frequency representation. J. Comp. Physiol. - A Sensory, Neural, Behav. Physiol. 181, 559–571 (1997).

45. Hackett, T. A., Barkat, T. R., O’Brien, B. M. J., Hensch, T. K. & Polley, D. B. Linking topography to tonotopy in the mouse auditory thalamocortical circuit. J. Neurosci. (2011).

46. Hu, R., Jin, S., He, X., Xu, F. & Hu, J. Whole-brain monosynaptic afferent inputs to basal forebrain cholinergic system. Front. Neuroanat. 10, 1–10 (2016).

47. Froemke, R. C. et al. Long-term modification of cortical synapses improves sensory perception. Nat. Neurosci. (2013).

48. Yerkes, R. M. & Dodson, J. D. The relation of strength of stimulus to rapidity of habit-formation. J. Comp. Neurol. Psychol. (1908).

49. Dana, H. et al. Sensitive red protein calcium indicators for imaging neural activity. Elife (2016).

50. Pachitariu, M. et al. Suite2p: beyond 10,000 neurons with standard two-photon microscopy. bioRxiv (2016).

